# A novel ex vivo bovine corneal infection and clearance model for Neisseria gonorrhoeae, Staphylococcus aureus, and Pseudomonas aeruginosa

**DOI:** 10.1101/2023.07.05.547903

**Authors:** Faith Uche Ukachukwu, Raid Alany, Lori A.S. Snyder

## Abstract

Ocular infections caused by bacterial pathogens may damage the cornea and rapidly progress to permanent blindness. Topical application of an ophthalmic formulation is often used to treat corneal infections. The animal models used in many preclinical studies frequently involve expensive *in vivo* experiments that compromises the corneal epithelium to mimic real life conditions such as during contact lens wear, but these do not consider other instances where infection occurs in intact corneas such as in ophthalmia neonatorum. To develop an *ex vivo* model of infection, bovine eyes from human food chain waste were processed and the corneas inoculated with *Neisseria gonorrhoeae* strain NCCP 11945*, Staphylococcus aureus* strain 6571, and *Pseudomonas aeruginosa* strain ATCC 15442 for 1 hour, 4 hours, and 6 hours, respectively. Inoculation included intact bovine corneas and those compromised with scalpel, needle, and blot methods. Recovery of *N. gonorrhoeae*, *S. aureus,* and *P. aeruginosa* colonies demonstrated that infection of bovine corneas was achieved with intact and compromised corneas using this model. In addition, corneas inoculated with the bacteria were treated with a suitable antibiotic, demonstrating clearance of the bacterial infections with at least 5 log_10_ reduction. This model is appropriate for both establishing infection and testing the ability of antimicrobial agents to clear bacterial eye infections. The bovine *ex vivo* model is reliable, cost-effective, suitable for different bacteria species, and reduces the need for further animal exploitation in laboratory research.

**Author Summary:** Bacterial pathogens such as *Neisseria gonorrhoeae*, *Staphylococcus aureus,* and *Pseudomonas aeruginosa* infect the eyes, damage the clear transparent cornea and may eventually cause blindness. Several *in vivo* animal models that have been used to investigate corneal infections in preclinical studies involve compromising the integrity of the corneal epithelium, which predisposes the eye to infection and simulates conditions of corneal abrasion suggested to be seen during contact lens wear. However, corneal infection in infants during ophthalmia neonatorum occurs with intact corneal epithelium and as such may not be explained by abrasion simulating models. Also, *in vivo* experiments are expensive, involve invasive corneal procedures despite efforts at ethical compliance, and may be time consuming. Reliable models that are quicker, cost effective, cause less (or no) discomfort to animals, and simulate a wide range of corneal infection scenarios need to be explored. Here, we demonstrate the use of a novel *ex vivo* bovine eye model to establish bacterial infection of the cornea, with and without compromising the corneal epithelium, and clearance of the infection with selected antimicrobial agents. The optimisation of the *ex vivo* bovine corneal infection model may serve as a bridge between *in vitro* and *in vivo* models of corneal infection.

## 1.0 Introduction

The cornea is an avascular ocular tissue that contributes to vision as a refractive medium for light entering the eye and provides protection to the ocular system against invasion by microbial pathogens [1,2]. Ocular infections that damage the cornea and progress to blindness can occur from bacteria pathogens such as *Neisseria gonorrhoeae, Staphylococcus aureus*, and *Pseudomonas aeruginosa* [3–7]. *N. gonorrhoeae*, responsible for the sexually transmitted infection gonorrhoea, also causes eye infections that commonly affect infants [8–10]. *S. aureus* and *P. aeruginosa*, associated with nosocomial infections and infections of the skin, lungs, ear, and heart in immunocompromised individuals, can also cause ocular infections at any stage in life [11–14]. Bacterial pathogens colonise and infect the eyes using mechanisms such as corneal adhesion and infiltration, traversal of the corneal layers, and stromal destruction [4]. The pathogenicity and virulence of these bacteria in corneal infections, potential therapeutic interventions, pharmacokinetics of ophthalmic formulations, and methods of drug delivery to the eye can be investigated to guide clinical management of these cases [15–18].

Human corneas are not routinely available for use in research because they are likely to be reserved for corneal transplant [19–20]. This therefore necessitates the development of non-human corneal models that closely represent conditions of the human corneal tissue. There are several ocular infection models, such as *in vitro* cell culture [21–23], *in vivo* animal models [5,24–27], and *ex vivo* animal models,[20, 26, 28–31] that can be used to understand the mechanisms of ocular infections. Currently, animal models often used in preclinical studies are *in vivo* models that involve use of large numbers of live animals that can be quite expensive and time consuming. These experiments may also utilize invasive corneal procedures that cause discomfort to the animals despite strict ethical guidelines and use of anaesthesia [20, 32]. In several animal models, to establish infection the corneal epithelium is usually compromised by scratching it with a needle [22,33,34] or scalpel [20, 30], or the corneal stroma is injected with the bacteria [20,30,35].

The susceptibility of the cornea seen with these invasive approaches is thought to be facilitated by the breach of the corneal integrity and is supported by similar conditions that may be observed in contact lens wear, such as corneal surface abrasion [20], or situations in which the unfavourable corneal conditions created by interaction of the contact lens with the tear fluid predisposes the cornea to infection [5]. However, in other conditions, corneal damage may occur following bacterial infection without prior compromise of the corneal epithelium. For instance, pathogenic microbial colonization of the neonatal cornea such as with *N. gonorrhoeae in utero* via antepartum infection of the placenta [36], ruptured placenta membrane that results in ascension of the bacteria in caesarean section [37–39], or during passage through the birth canal of an infected mother [40–42]. Despite an intact corneal epithelium, introduction of *N. gonorrhoeae* to the ocular surface may result in gonococcal conjunctivitis in new-borns and adults (via autoinoculation of infected genital secretions), which is capable of causing corneal ulceration, scarring, and blindness within 24 hours [9,43–49]. Thus, ocular models are needed that both closely represent this condition and enable susceptibility assays for research and development of novel treatments [50].

Animals such as rats [61–65], rabbits [20,61,66–68], pigs [69, 70–72], mice [26, 27,61, 73, 74], dogs [31,75,-78]), and goats [18, 79, 80] have been used as *in vivo* models of infection in previous studies. Alternative methods that are viable and eliminate pain to animals have been encouraged [51–60].

*Ex vivo* ocular models that are used in corneal studies usually involve animal corneal tissues that are excised and prepared for laboratory investigations [19]. While a few *ex vivo* models have been used previously [20,26,28–31, 81–84], there is paucity of studies that utilize an *ex vivo* model to investigate corneal infection caused by several bacterial pathogens to demonstrate suitability for different organisms. In addition, an *ex vivo* model that utilizes corneal tissues that would otherwise be waste products from human food production, rather than those excised from animals specifically for laboratory investigations, aligns with the 3Rs principles of humane experimental technique (Replacement, Reduction, and Refinement of use of live animals) [53,85–90].

Here, an *ex vivo* bovine eye model that is simple, reproducible, cost effective, and time saving is used to investigate bacterial infection of the cornea caused by *N. gonorrhoeae, S. aureus*, and *P. aeruginosa.* Moreover, the topical route of ophthalmic drug application is one of the mainstays of corneal infection management due to simplicity of administration, potential to establish therapeutic concentrations at the target corneal site, and ability to achieve localised therapeutic and/or adverse action [19]. Therefore, the utility of the bovine corneal model to investigate standard antibiotics that might be used topically for treatment of gonococcal, staphylococcal, and pseudomonad eye infections is also explored. This *ex vivo* bovine eye model explores corneal infection that occurs with intact and compromised corneal epithelium as a potential bridge between *in vitro* and *in vivo* models of ocular infection. There is promising potential to use this model to explore bacterial ocular infections and to develop novel antimicrobials.

## 2.0 Materials and methods

### 2.1 Bacterial strains and culture

The bacterial strains used were *N. gonorrhoeae* strain NCCP 11945, *S. aureus* strain 6571, and *P. aeruginosa* strain ATCC 15442. *N. gonorrhoeae* was cultivated on GC agar medium (Oxoid) including the Kellogg supplements [91]. The Kellogg’s glucose supplement was made by dissolving 20 g of D-glucose in 70 ml distilled water on a stirring hot plate. Once the glucose is dissolved and the solution cooled to room temperature, 0.5 g of L-glutamine and 0.001 g of co-carboxylase (thymine pyrophosphate) were added; the final volume was taken up to 100 ml, and the supplement was filter sterilized. Kellogg’s iron supplement is 0.5% w/v ferric nitrate (Fe(NO_3_)_3_ in 10 ml distilled water, sterilized by filtration. Both supplements were stored at 4°C. *N. gonorrhoeae* agar plates were incubated for 24 to 48 hours in a CO_2_ (5% v/v) incubator at 37°C. To prepare the gonococci for inoculation of the bovine corneas, piliated (P+) colonies of *N. gonorrhoeae* were selected based on colony appearance and harvested into GC broth with the Kellogg’s iron supplement. *S. aureus* and *P. aeruginosa* were grown on nutrient agar (Fisher Scientific) and incubated for 24 hours at 37°C. In preparation for bovine corneal *ex vivo* infection, colonies of these bacteria were harvested into nutrient broth. For inoculation, bacterial resuspensions of *N. gonorrhoeae* in GC broth were adjusted to OD_600_ 0.5 (about 10^8^ colony forming units per millilitre (cfu/ml)), while *S. aureus* and *P. aeruginosa* were adjusted in nutrient broth to OD_600_ 0.4 (approximately 10^9^ cfu/ml) and OD_600_ 0.3 (around 10^9^ cfu/ml), respectively, and then adjusted to 0.5 McFarland turbidity standard (∼10^8^ cfu/ml).

### 2.2 Antimicrobial agents

Ceftriaxone and ciprofloxacin (Merck) stocks were made at 1000 µg/ml in distilled water. Phosphate buffered saline (PBS) was used as a negative control for corneal clearance infection experiments.

### 2.3 Antimicrobial susceptibility testing

Broth microdilution was used to determine the minimum inhibitory concentration (MIC) of the antibiotics in accordance with the Clinical Laboratory Standards Institute (CSLI) guidance [92] with some modifications, as previously [93,94]. In brief, doubling dilutions of stock antibiotic solutions were prepared in 96 well plates in triplicate wells to a volume of 100 µl using double strength broth. Ceftriaxone broth microdilutions used double strength GC broth to determine the MIC for *N. gonorrhoeae* strain NCCP11945. Ciprofloxacin in double strength nutrient broth was used to assess the MICs of *S. aureus* and *P. aeruginosa.* Addition of the bacteria to each well resulted in single strength broth. The concentrations of the antibiotics in the wells ranged from 500 µg/ml to 5.96 x10^-5^ µg/ml (24 dilutions) and three independent experiments were done (n=3) on different days to generate biological replicates.

To prepare the bacterial inoculum, *N. gonorrhoeae* was suspended in GC broth and measured spectrophotometrically at OD_600_ 0.5, which is roughly equivalent to 1.5 × 10^8^ cfu/ml. For *S. aureus* and *P. aeruginosa*, nutrient broth was used and the bacterial suspensions were measured at OD_600_ 0.4 (approximately 1.5 × 10^9^ cfu/ml) and OD_600_ 0.3 (approximately 1.5 × 10^9^ cfu/ml), respectively. To each well of the 96 well plate was added 100 µl of freshly prepared bacteria suspension to achieve a final volume of 200 µl in the wells and bacterial concentrations of about 5 × 10^6^ cfu/ml for *N. gonorrhoeae* and 5 × 10^5^ cfu/ml for *S. aureus* and *P. aeruginosa.* Viable counts of the bacterial concentrations were confirmed by serially diluting 10 µl of the suspension or microtiter plate volume and enumerating colonies on the respective agar plates.

The controls used for the experiment include a broth contamination control well containing broth only, an antimicrobial control containing only broth and antibiotics to assess any impact on well appearance due to the antimicrobial, a growth control with broth and bacteria, but no antibiotics to evaluate bacteria growth. Of these three, the latter is used for comparison of growth in the MIC determining wells and the former two are used for comparison of wells that have not produced growth.

The 96 well plates were incubated at 37°C for 24 hours; *N. gonorrhoeae* cultures used a 5% CO_2_ incubator. Absorbances for each well were read at OD_600_ by a microplate reader infinite series spectrophotometer (Tecan). The lowest antibiotic concentration that after 24 hours incubation had an optical density in line with the antimicrobial control was determined to be the MIC. Following spectrophotometric assessment, 10 µl from each well of the 96 well plates was seeded on to the respective agar plates and then incubated under standard conditions for 24 hours. The lowest concentration of antibiotics for which there was no growth of bacteria on the agar plate was considered as the minimum bactericidal concentration (MBC).

### 2.4 Bovine eye model of corneal infection

#### 2.4.1 Preparation and infection of uncompromised corneas

*Ex vivo* bovine corneal infection was conducted with slight modification from that described previously [95]. Bovine eyes were collected from an abattoir (ABP Guildford, Surrey, UK), where they are the by-product of human food production. To reduce risk of desiccation and contamination, excised bovine eyes were transported in cold PBS containing Pen-Strep antibiotics (Fisher Scientific) and antimycotic amphotericin B (Fisher Scientific) (PBS+A&A). Upon arrival, the bovine eyes were either immediately processed or stored in fresh PBS+A&A at 4°C and used within 5 days.

To process the eyes, corneas were examined for gross defects, such as scratches or lacerations (Fig 1A). Corneas that were suitable were excised from the eye using scissors sterilized in ethanol, with the assistance of forceps (Fig 1B). The excised corneas were then rinsed first in PBS+A&A, then in sterile PBS, and placed epithelial side down (exterior surface) on sterile spot plates to help hold the shape for adding the scaffold medium. Molten scaffold medium was prepared by heating a mixture of agarose (0.2 g) and RPMI medium (20 ml) (Fisher Scientific); this was added on the endothelial side (interior) of the corneas and allowed to cool to retain the corneal shape (Fig 1C).

**Fig 1:**
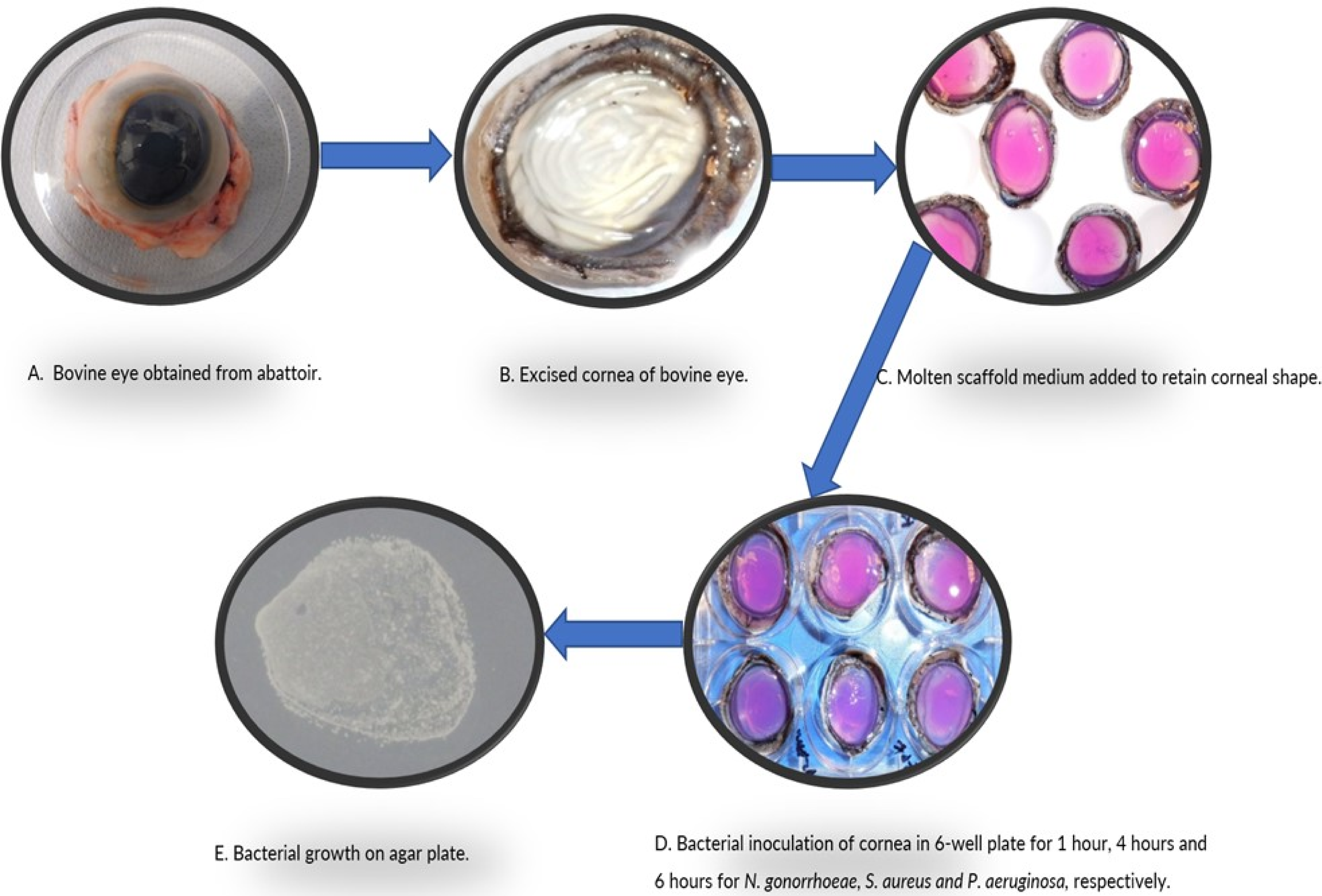
Stages of infecting uncompromised cornea in the *ex vivo* bovine eye model of infection. Infection of bovine corneas with *N. gonorrhoeae, S. aureus*, and *P. aeruginosa* involved various stages. (A) Bovine eyes were collected from a local abattoir. (B) After gross examination for any defects, bovine eyes with good corneas were excised using sterile forceps and scissors. (C) Molten scaffold medium made by heating agarose (0.2g) in RPMI (20ml) was added to the endothelial side (back surface) to maintain the corneal shape. (D) The corneal epithelium (front surface) was placed on 200 µl of bacterial suspension in 6-well plate and incubated (CO_2_ (5% v/v)) at 37°C for 1 hour, 4 hours, and 6 hours for *N. gonorrhoeae, S. aureus,* and *P. aeruginosa,* respectively. (E) Infected cornea (epithelial side) was rinsed with PBS and pressed onto an agar surface (GC agar for *N. gonorrhoeae*, nutrient agar for *S. aureus* and *P. aeruginosa*). An imprint of bacterial growth was found after incubation of agar plates at 37°C for 24 hours. Triplicate experiments were done on the same day.

The corneas were not further cultured in any growth media before bacterial inoculation was done. The corneas were then placed epithelial side down in 6-well plates containing 200 µl of inoculum of each bacterium, prepared as for the microtiter plate assays, and kept at 37°C in a CO_2_ (5% v/v) incubator for infection times of 1 hour, 4 hours, and 6 hours (Fig 1D). Experiments were conducted in three replicates on the same day to generate technical replicates.

#### 2.4.2 Infection of compromised corneas

Apart from infecting the corneas with intact epithelium as described above, four additional methods that involved compromising the corneal epithelium were used in this study (Fig 2). Compromised corneas were infected with *S. aureus* and *P. aeruginosa* as for 2.4.1. Triplicate results were obtained for each.

**Fig 2:**
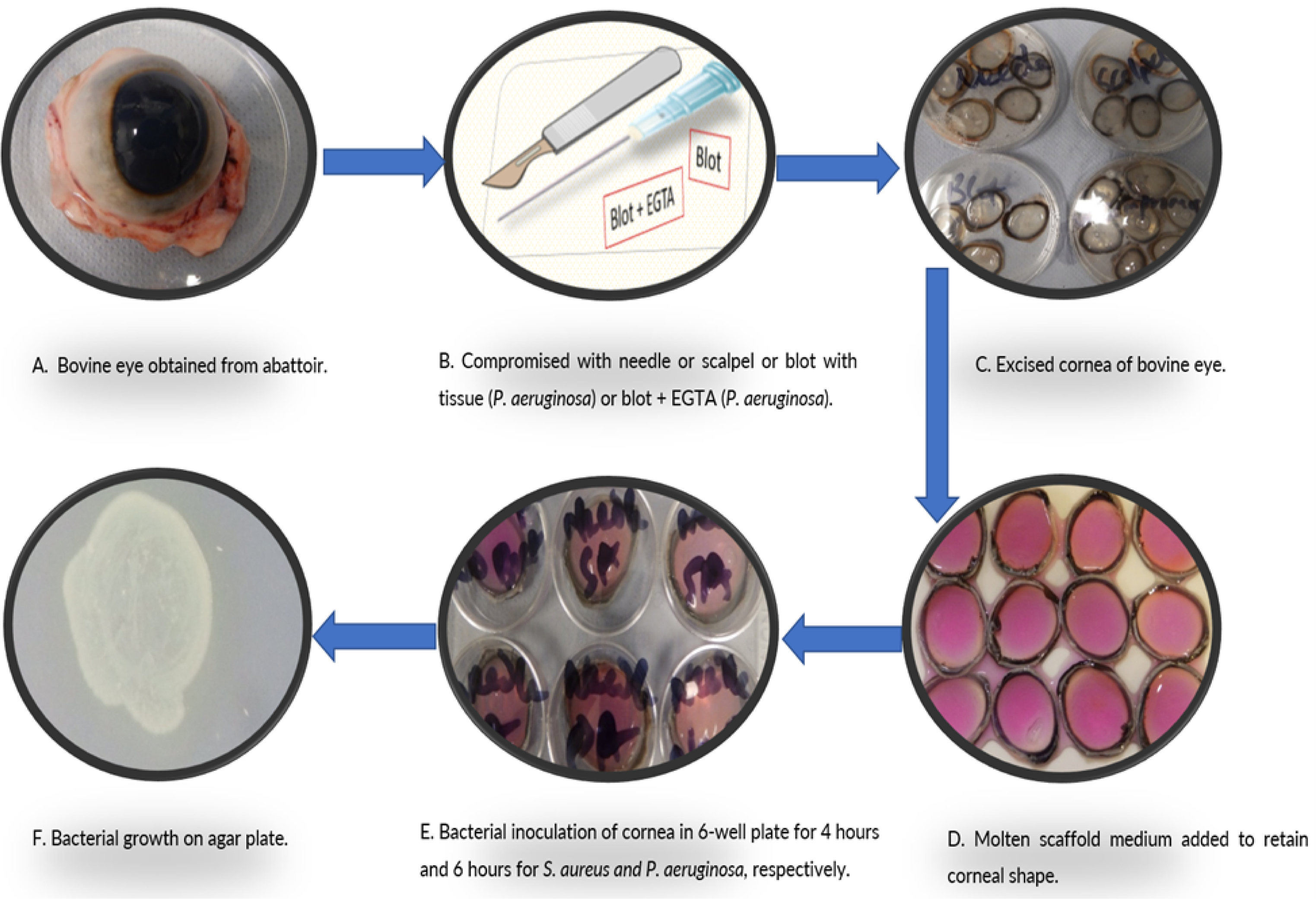
Stages of infecting compromised cornea in *ex vivo* bovine eye model of infection. The process of compromising bovine corneas and infecting them with *S. aureus* and *P. aeruginosa* is similar to the method for uncompromised cornea, but with different initial steps. (A) Bovine eyes were obtained from a local slaughterhouse. (B) After inspection and before corneas were excised, bovine eyes with healthy corneas were compromised by using a 25-guage needle or scalpel to induce full thickness abrasion by making three vertical and horizontal incisions that overlap on corneal epithelium. For other corneas, the corneal epithelium was blotted 2 to 3 times with paper towel to compromise the cornea. Blotted corneas were either used following this treatment or further treated after excision with 400 µl of EGTA (100mM in PBS) and incubated (CO_2_ 5% v/v) at 37°C for 1 hour before infection (not shown). (C) Corneas were excised with sterile forceps and scissors. (D) Corneal shape was maintained by adding a molten scaffold medium (heated agarose 0.2 g in RPMI 20 ml) to the internal surface (endothelial side). (E) Corneas were placed epithelial side (front surface) down in 6 well plates containing 200 µl of bacteria suspension and kept in CO_2_ (5% v/v) incubator at 37°C for 4 hours and 6 hours for *S. aureus* and *P. aeruginosa*, respectively. (F) After rinsing with PBS, the epithelial side of infected cornea was pressed onto a nutrient agar surface to obtain an imprint of bacterial growth following 24 hours incubation of agar plates at 37°C. These methods involving compromised corneas were not done for *N. gonorrhoeae*. The Blot and Blot + EGTA method were only done for *P. aeruginosa*. Three replicates of the experiment were done on the same day.

##### 2.4.2.1 Needle method

This technique was adapted from work done previously [22,96]. In brief, before the corneas were excised, a 25-guage needle (Fisher Scientific) was used to make three 1-mm parallel lines running vertically and across horizontally overlapping on the epithelium of the cornea to induce full thickness abrasion. After the corneas were excised and rinsed with PBS, the scaffold medium was added to the endothelial side of the cornea and then inoculated with bacteria as described above (section 2.4.1).

##### 2.4.2.2 Scalpel method

This method was similar to that previously described [20]. Just as with the needle, a scalpel was used to induce full thickness abrasion of the corneal epithelium by making three vertical incisions and three horizontal incisions overlapping on the cornea. Infection of the cornea was done in similar fashion as already described (section 2.4.1).

##### 2.4.2.3 Blot method

This was adapted from previously described work [26]. Briefly, a sterile paper towel was used to blot the corneal epithelium of the bovine eye 2 to 3 times before the cornea was excised. Then the corneal infection was done as earlier described (section 2.4.1). The blot method was used with *P. aeruginosa* only.

##### 2.4.2.4 Blot and EGTA method

This approach was performed as previously [26]. Blotting of the corneal epithelium with a sterile paper towel 2 to 3 times was done as for the blot method (section 2.4.2.3). After excision and rinsing of the cornea with PBS, the cornea containing molten scaffold medium was placed epithelial side down in a 6 well-plate containing 400 µl of Ethylene glycol-O,O’-bis(2-aminoethyl)-N,N,N’,N’-tetraacetic acid (EGTA) (Fisher scientific) and was kept in a CO_2_ (5% v/v) incubator at 37°C for 1 hour. EGTA was prepared as 100mM in PBS. The corneas were then rinsed with PBS to remove the EGTA and inoculated with bacteria as described (section 2.4.1). The blot and EGTA method was used only with *P. aeruginosa*.

#### 2.4.3 Recovery of adherent bacteria

After incubating the inoculated corneas for the various infection hours, the epithelial side of the corneas were carefully rinsed with PBS to remove any non-adherent bacteria without dislodging the scaffold. The epithelial side of the corneas infected with *N. gonorrhoeae* were then imprinted on GC agar with Kellogg’s glucose and iron supplements. These agar plates were incubated at 37°C with CO_2_ (5% v/v). Corneas infected with *S. aureus* and *P. aeruginosa* were imprinted on nutrient agar and the plates were incubated at 37°C. After 24 hours, the bacteria transferred from the surface of the bovine corneas were recovered on the agar plates and assessed (Figs 1E and 2F).

#### 2.4.4 Recovery and quantification of internalised bacteria

To recover the bacteria that had been internalised by the corneal cells, the inoculated corneas were rinsed with PBS following incubation for the various infection hours. The epithelial cornea was placed on 100 µl of saponin solution (1% (w/v) saponin in PBS; Fisher Scientific) in 6-well plates and incubated for 10 minutes at 37°C in 5% (v/v) CO_2_. The epithelium was then scraped with a sterile scalpel to expose the inner corneal layers and rinsed in the saponin solution from the well of the 6-well plate. This 100 µl of saponin solution was then removed from the well via pipette, homogenised in 900 µl of appropriate broth (GC or nutrient), and serially diluted 1:10. In each case, 100 µl of the serial dilutions were plated in duplicate on appropriate agar plates. Plate counts of colonies that appeared on the agar plates were documented as internalised bacteria.

#### 2.4.5 Clearance of corneal infection with antimicrobial agent

One hour after inoculation of the corneas with bacteria, the corneal tissue was rinsed and placed on a petri dish with the epithelial side facing upwards. To assess the antimicrobial’s ability to clear the bacteria from the corneal surface, 500 µl of the test sample was added onto each cornea, left for a time suitable to test the treatment, and rinsed off with PBS. Adherent and internalised bacteria were recovered as colonies on agar plates after incubation (section 2.4.3 and 2.4.4). Ceftriaxone was tested on corneas infected with *N. gonorrhoeae* while ciprofloxacin was used to challenge *S. aureus* and *P. aeruginosa* infection of corneas. In ophthalmia neonatorum prophylaxis, the goal is to apply the antimicrobial prophylaxis within one hour of birth, thus the chosen time for inoculation and incubation of the corneas before treatment with the antimicrobial. For the negative control, PBS was applied to the infected corneas instead of an antibiotic. Tests were performed in triplicates.

## 3.0 Results

### 3.1 Corneal imprints and colonies of bacteria on agar plates

Corneal imprints of the bacteria showed recovery of *N. gonorrhoeae* that adhered to the surface of the cornea after 1 hour of infection (Fig 3A). *S. aureus* adhered to the corneal surface after infection for 4 hours (Figs 3C, 4A and 4C) and *P. aeruginosa* at 6 hours (Figs 3E, 4E, 4G, 4I and 4K). The different methods of establishing infection on uncompromised, intact corneas (Fig 3) and also compromised corneal epithelium using needle, scalpel, blot, and blot and EGTA (Fig 4), were demonstrated qualitatively via corneal print. In each case, bacteria were recovered on the agar plates that had adhered to the corneal surfaces.

**Fig 3:**
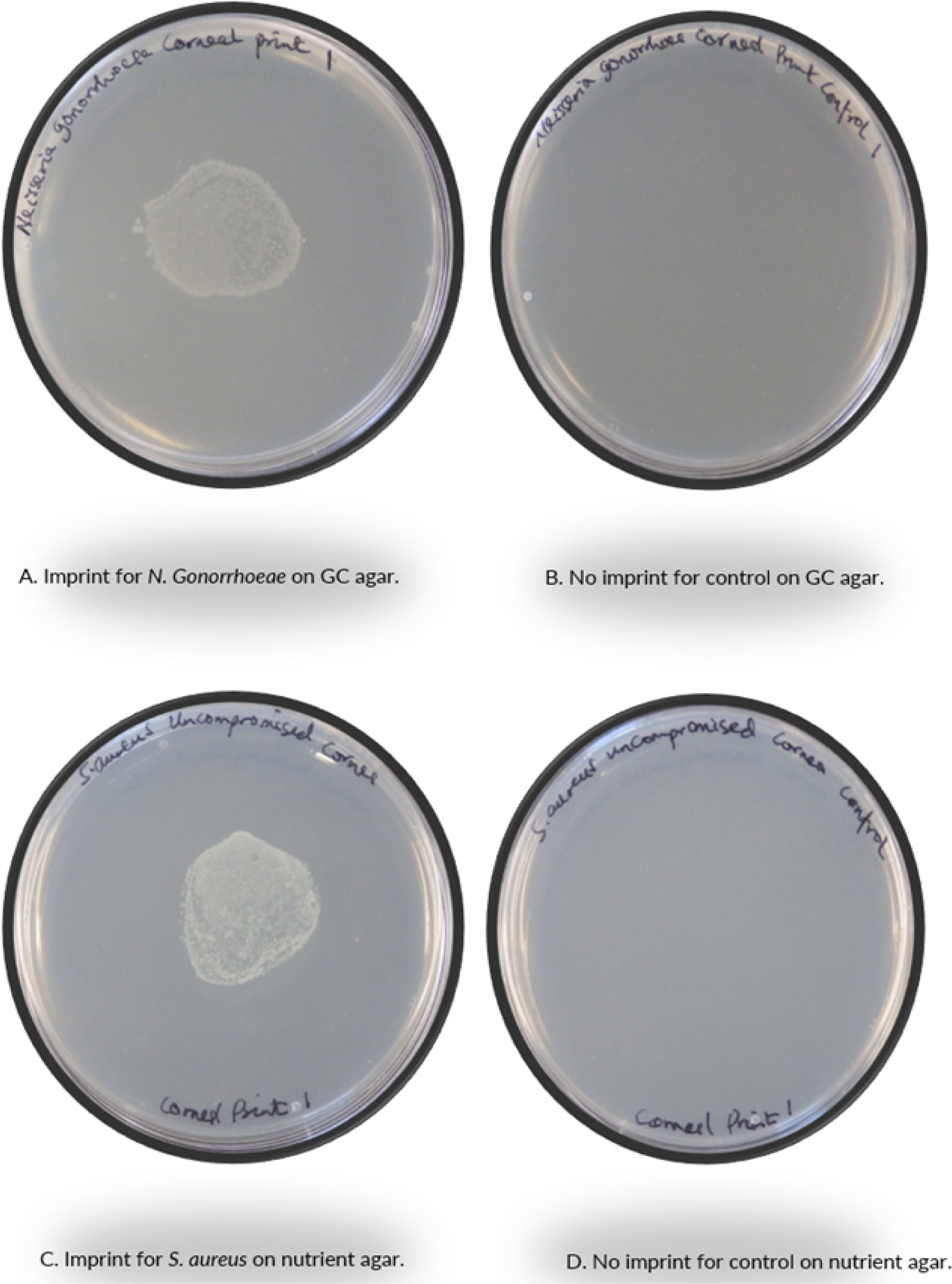
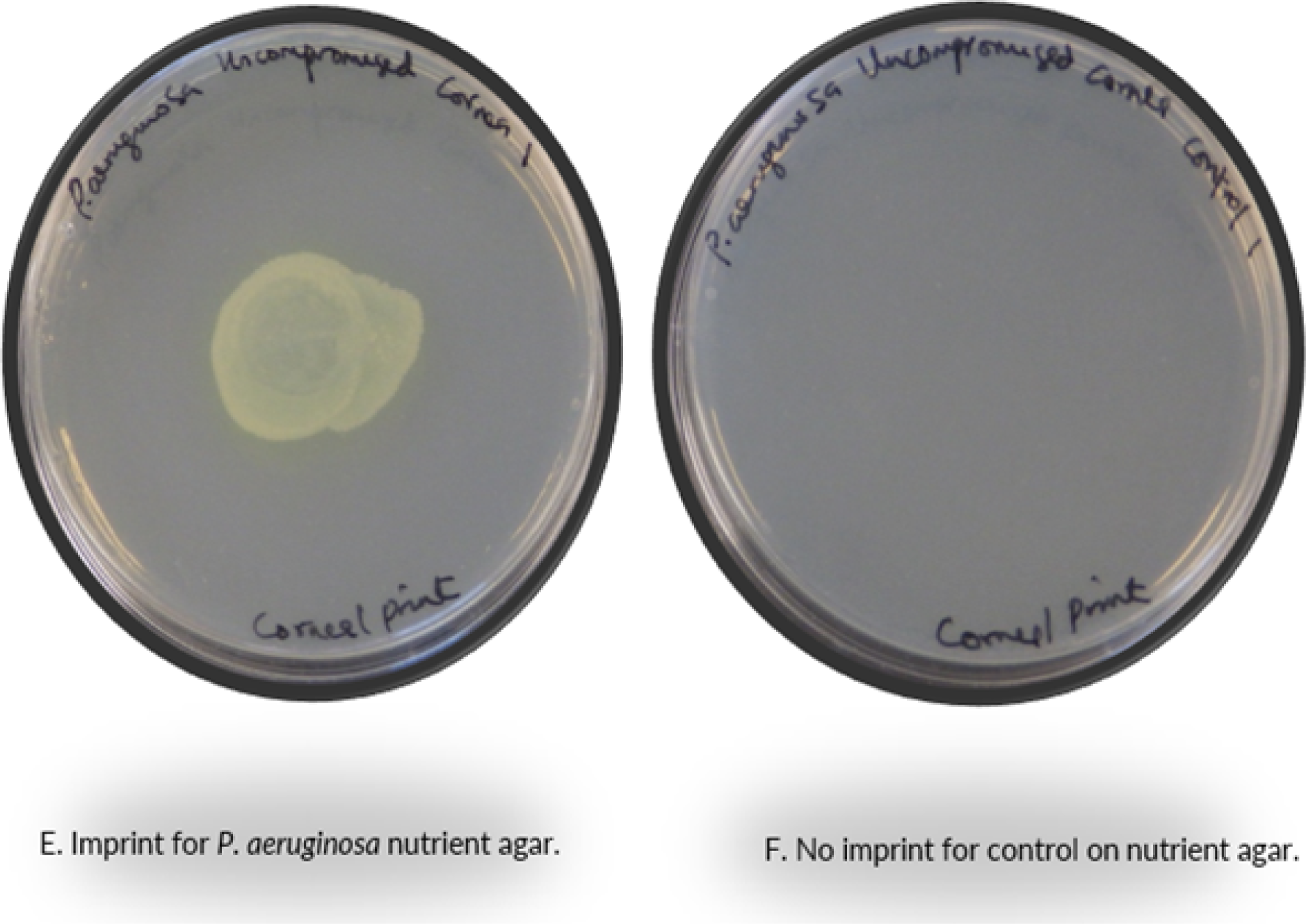
Epithelial adhered bacteria recovered from infected uncompromised corneas. Qualitative data for bacteria adherence is depicted as presence or absence of growth from corneal imprints. Bovine corneal inoculation was done with *N. gonorrhoeae*, *S. aureus*, and *P. aeruginosa*. Respective broths were used, and growth was sampled on respective agar plates. The agar plates were incubated for 24 hours at 37°C. There was recovery of bacterial colonies for corneas infected with *N. gonorrhoeae* (A), *S. aureus* (C), and *P. aeruginosa* (E), in contrast to the no growth on the respective controls (B, D, and F).

**Fig 4:**
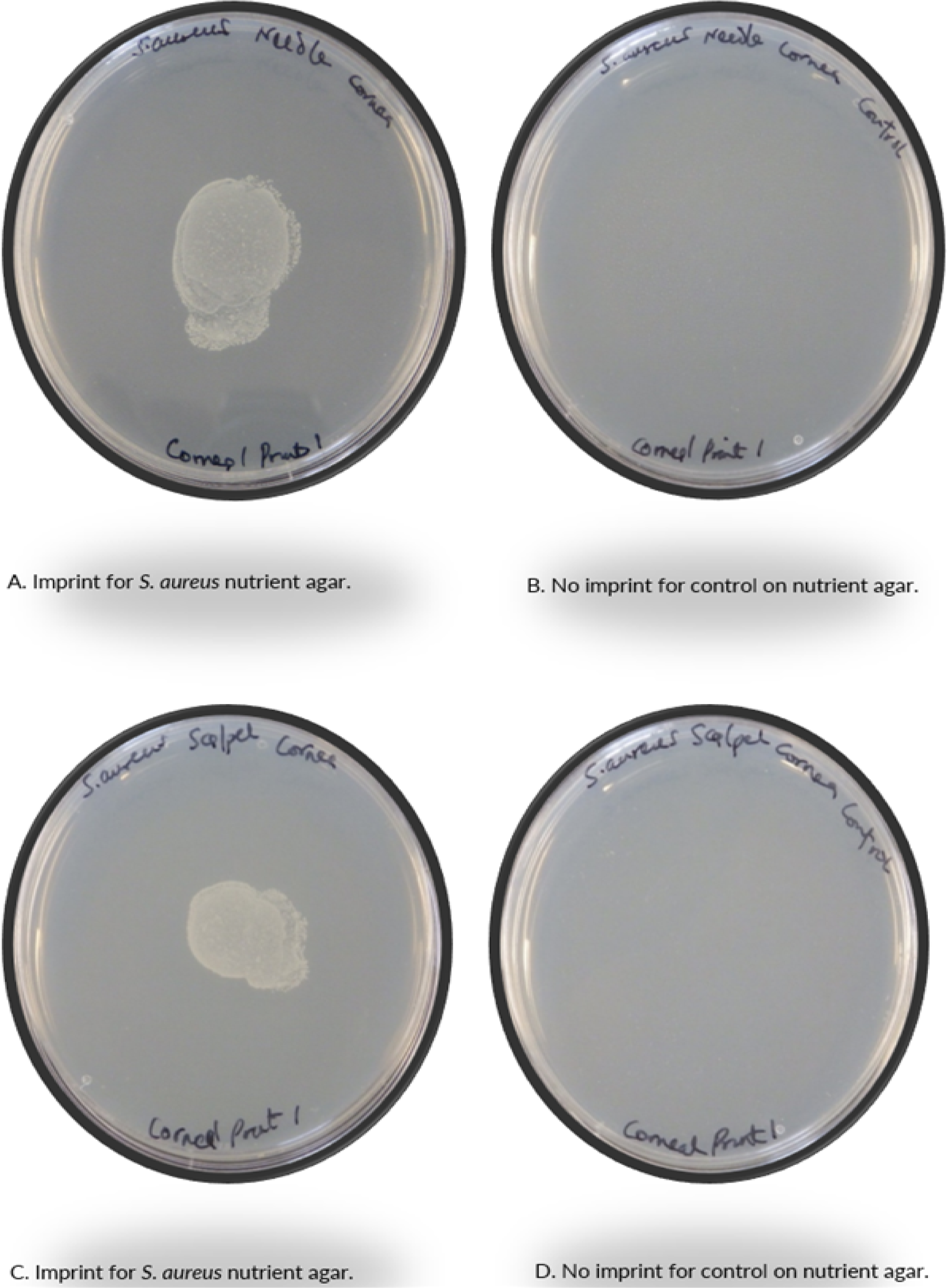
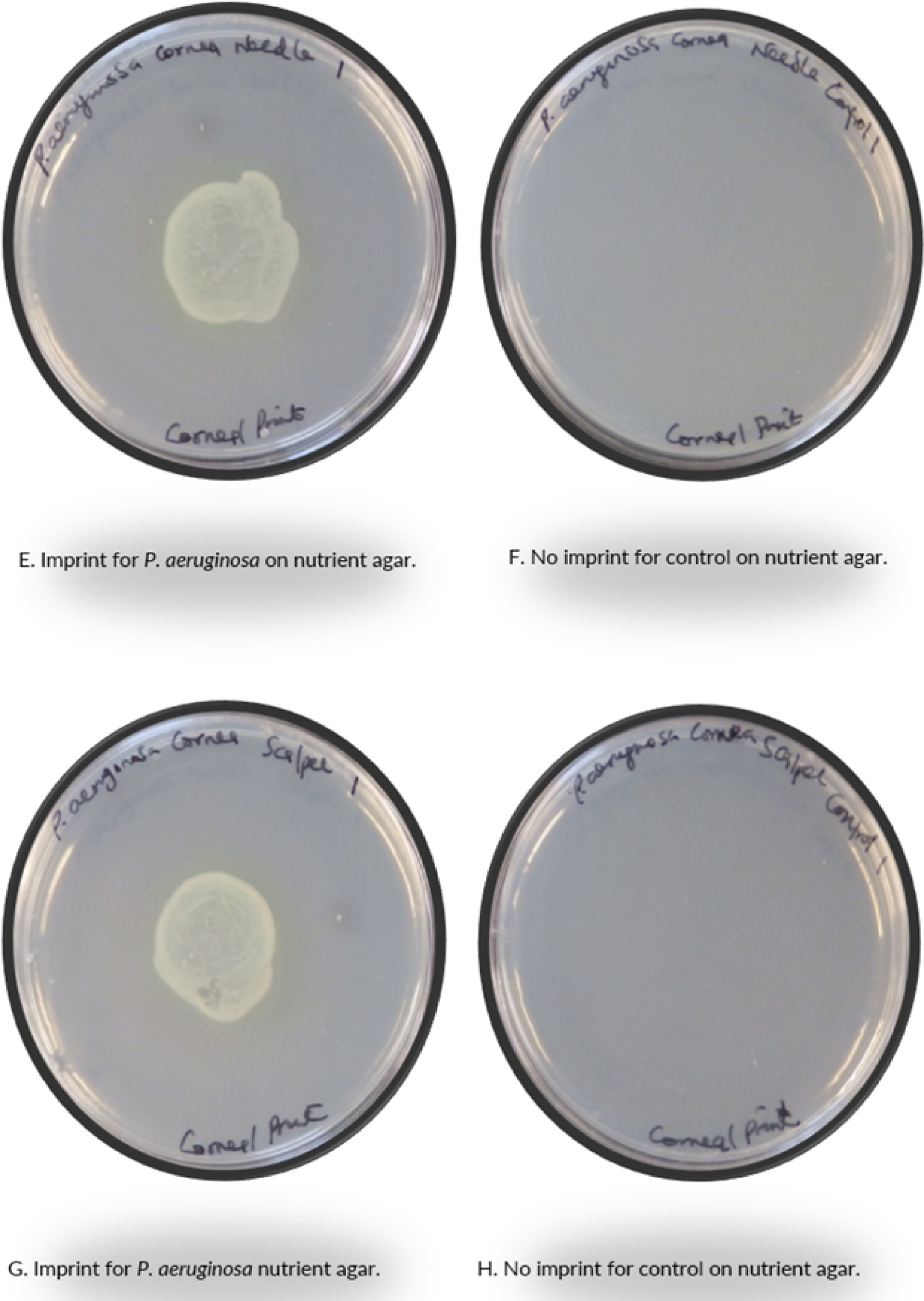
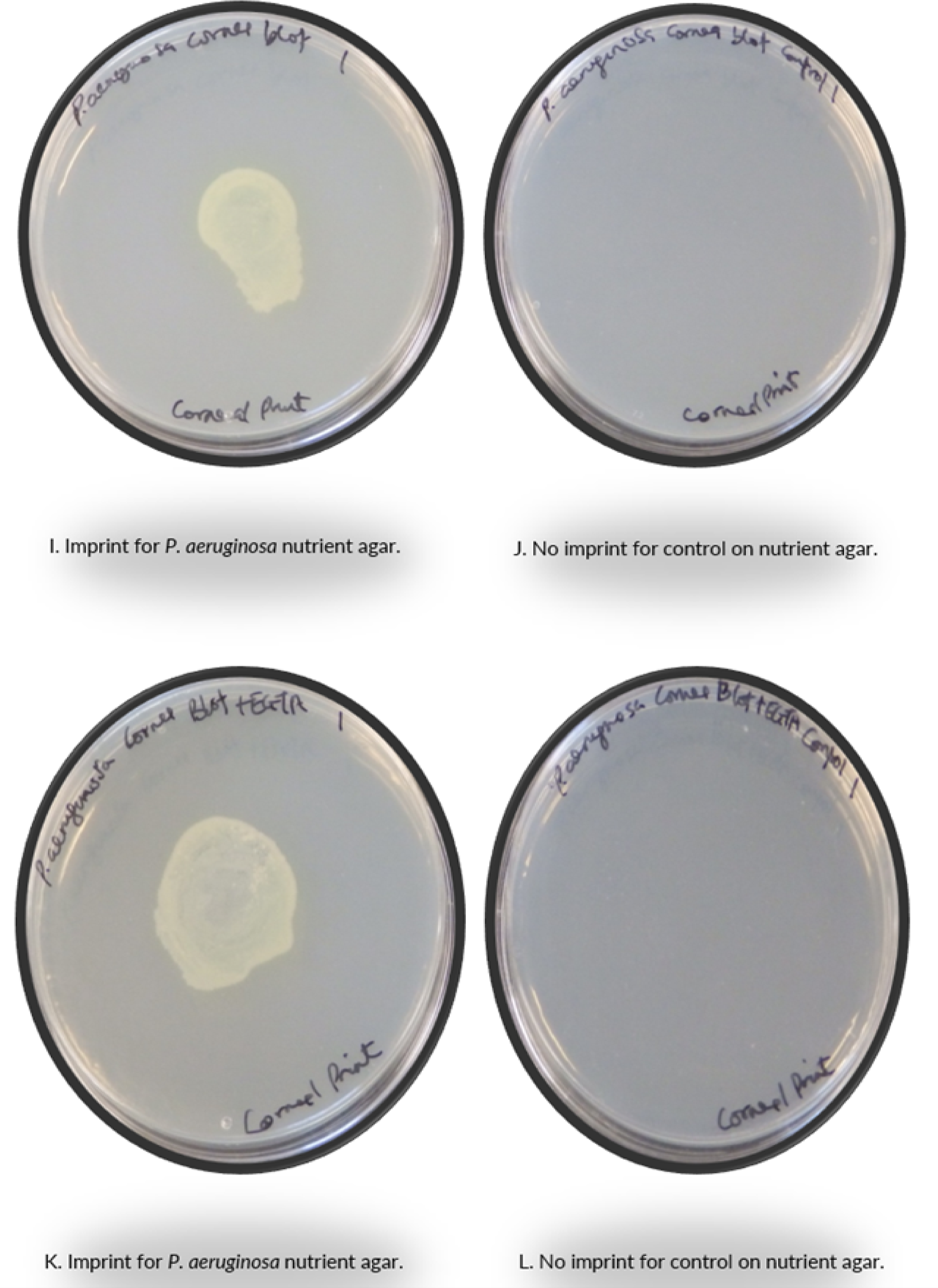
Adhered bacteria recovered from infecting compromised corneas. Qualitative results for bacteria adherence to corneas with compromised epithelium is shown as presence or absence of growth on corneal imprints. Corneal infection of compromised, excised bovine corneas was done with *S. aureus* and *P*. *aeruginosa*, sampled on nutrient agar plates incubated for 24 hours at 37°C. The bovine corneas were compromised by one of several methods: needle scratches (A, B, E, and F), scalpel cuts (C, D, G, and H), blots with paper towel (I and J), and blot + treatment with EGTA (K and L). Bacterial colonies were recovered on nutrient agar for corneas infected with bacteria (A, C, E, G, I, and K), unlike controls (B, D, F, H, J, and L).

Gonococci internalized by the excised bovine corneas were recovered 1 hour after infection (Figs 5A and 5C). Staphylococcal recovery was seen at 4 hours (figs 5E, 5G, 6A, 6C, 6E and 6G) and pseudomonad at 6 hours (figs 5I, 5K, 7A, 7C, 7E, 7G, 7I, 7K, 7M and 7O).

**Fig 5:**
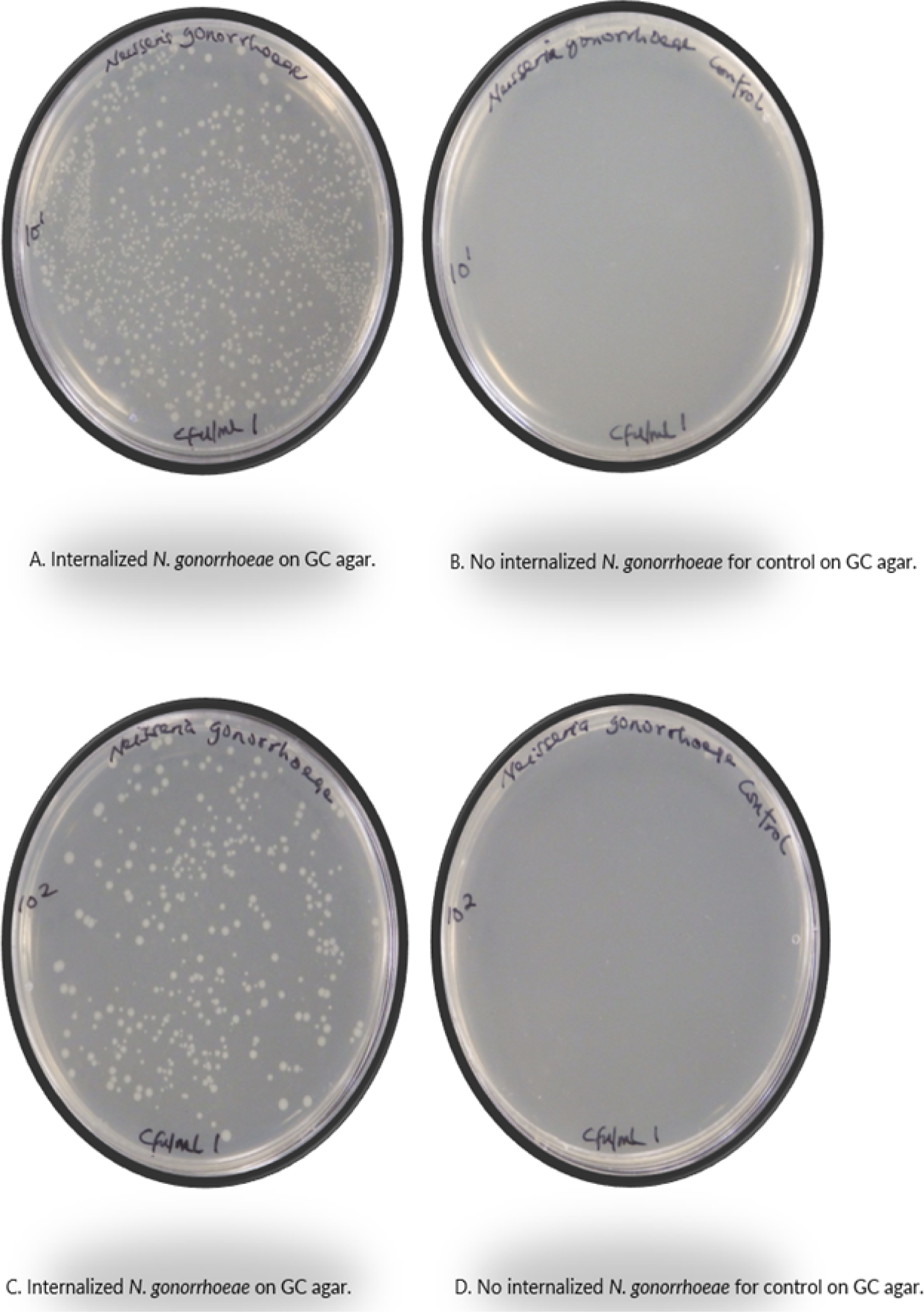
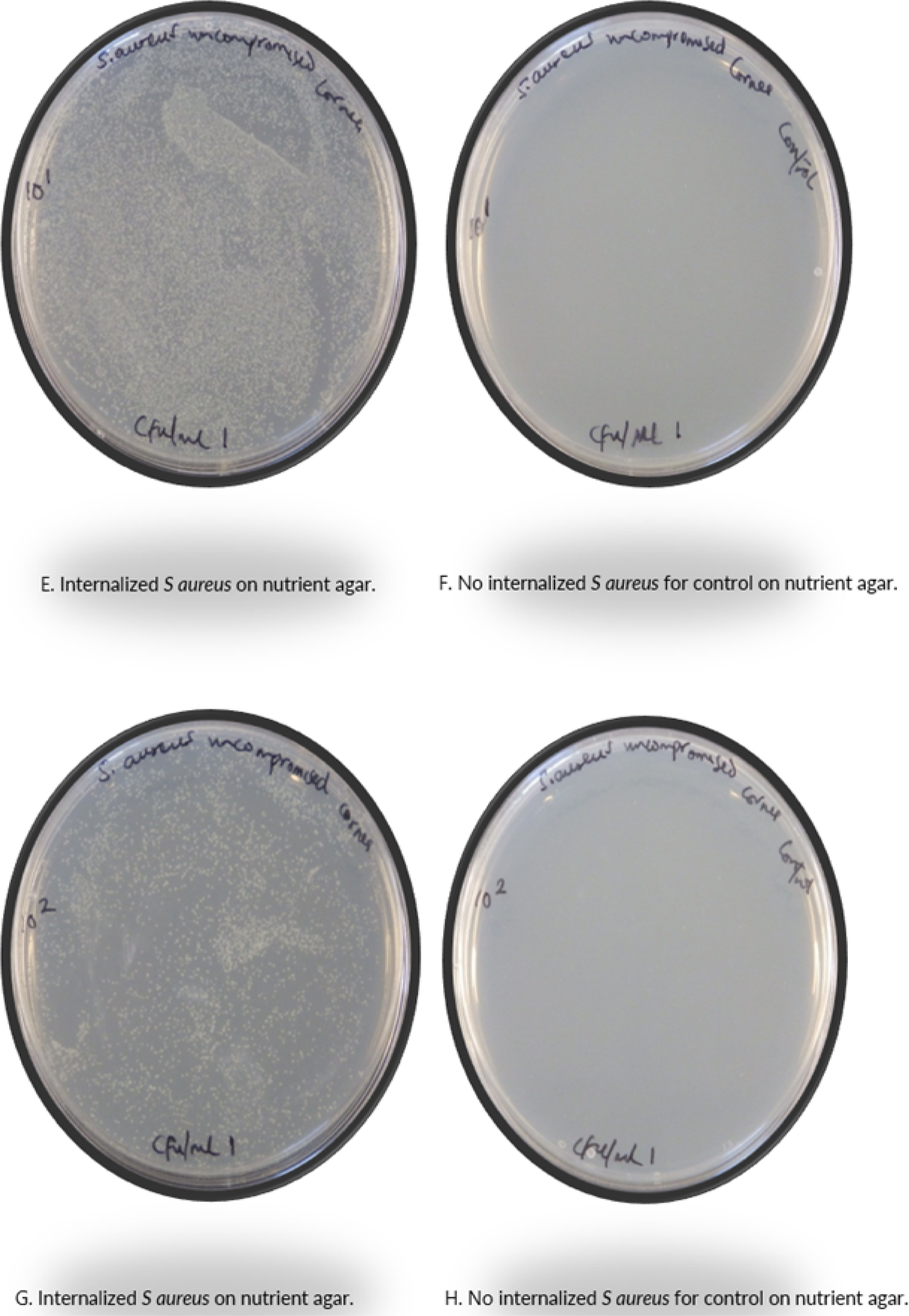
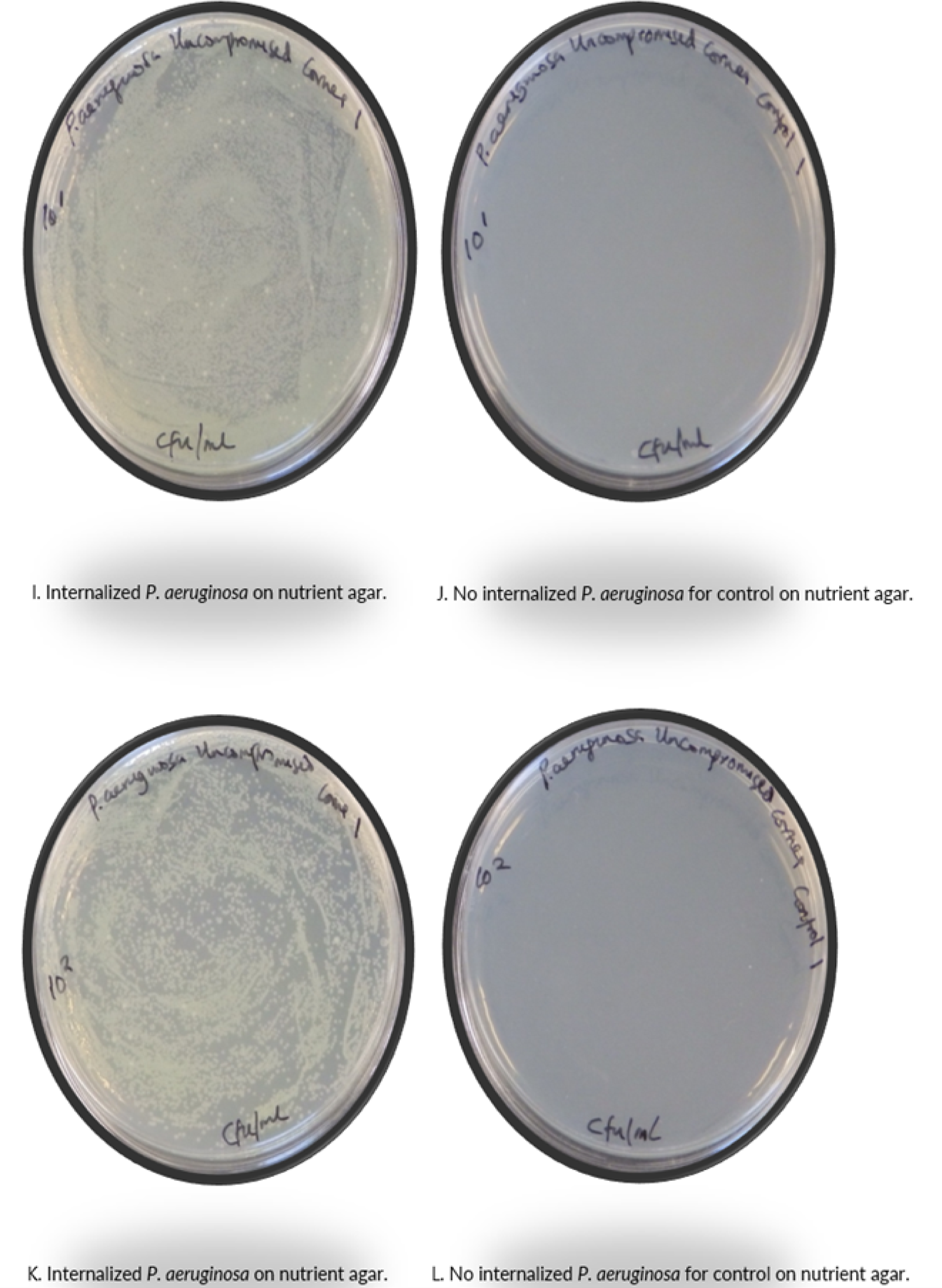
Internalized bacteria recovered from intact corneas. Corneal infection of bovine eyes was done with *N. gonorrhoeae*, *S. aureus,* and *P. aeruginosa* and internalized bacteria were recovered with 1% (w/v) saponin plated onto respective agar plates. The agar plates were incubated for 24 hours at 37°C. Colonies of *N. gonorrhoeae* were recovered on GC agar (A, C), and those of *S. aureus* (E, G) and *P. aeruginosa* (I, K) on nutrient agar plates. In contrast, there was no growth on the controls (B, D, F, H, J and L). Quantitative data for presence or absence of internalized bacteria was obtained from serial dilutions of up to 10^2^.

**Figure 6:**
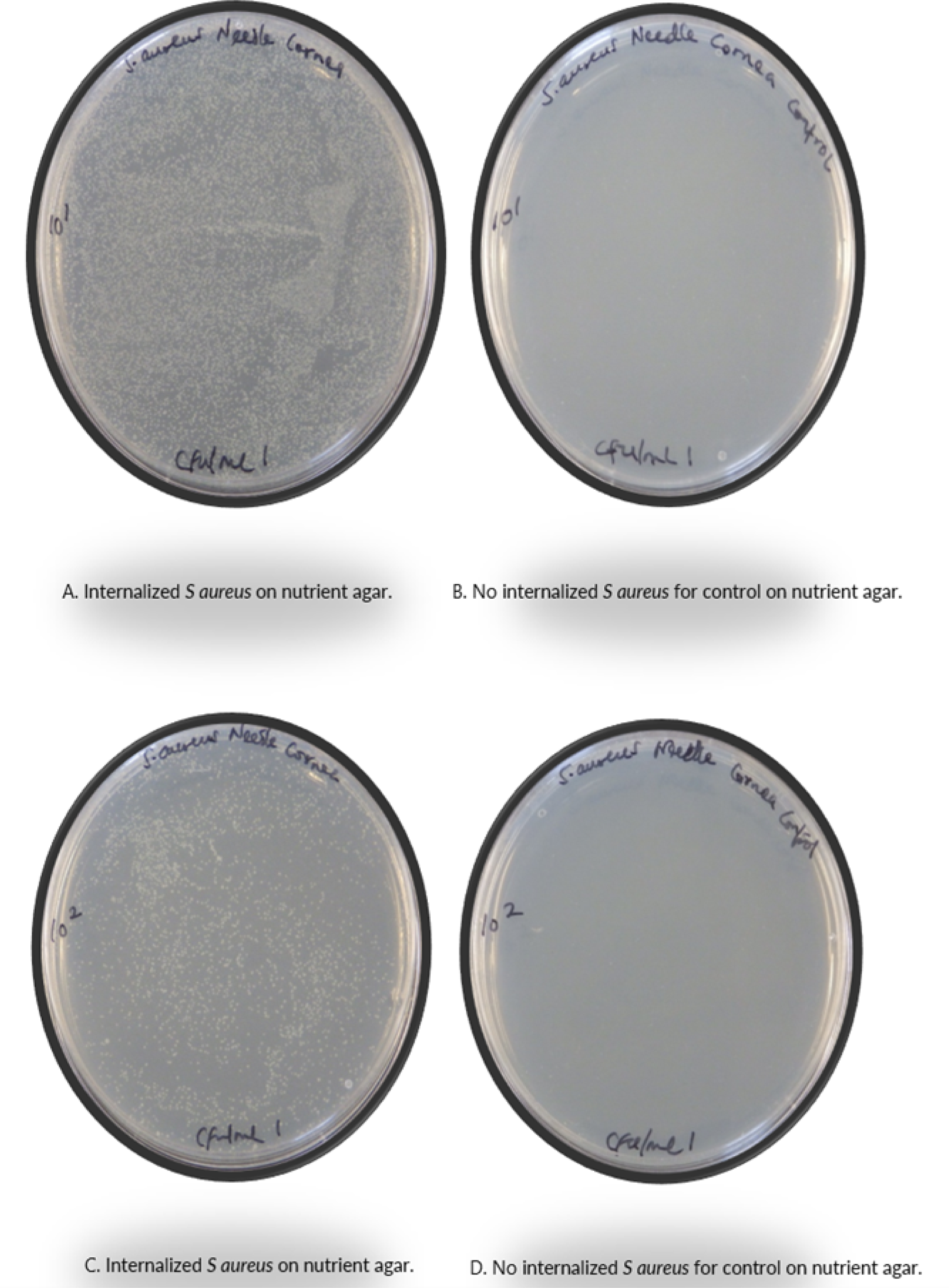
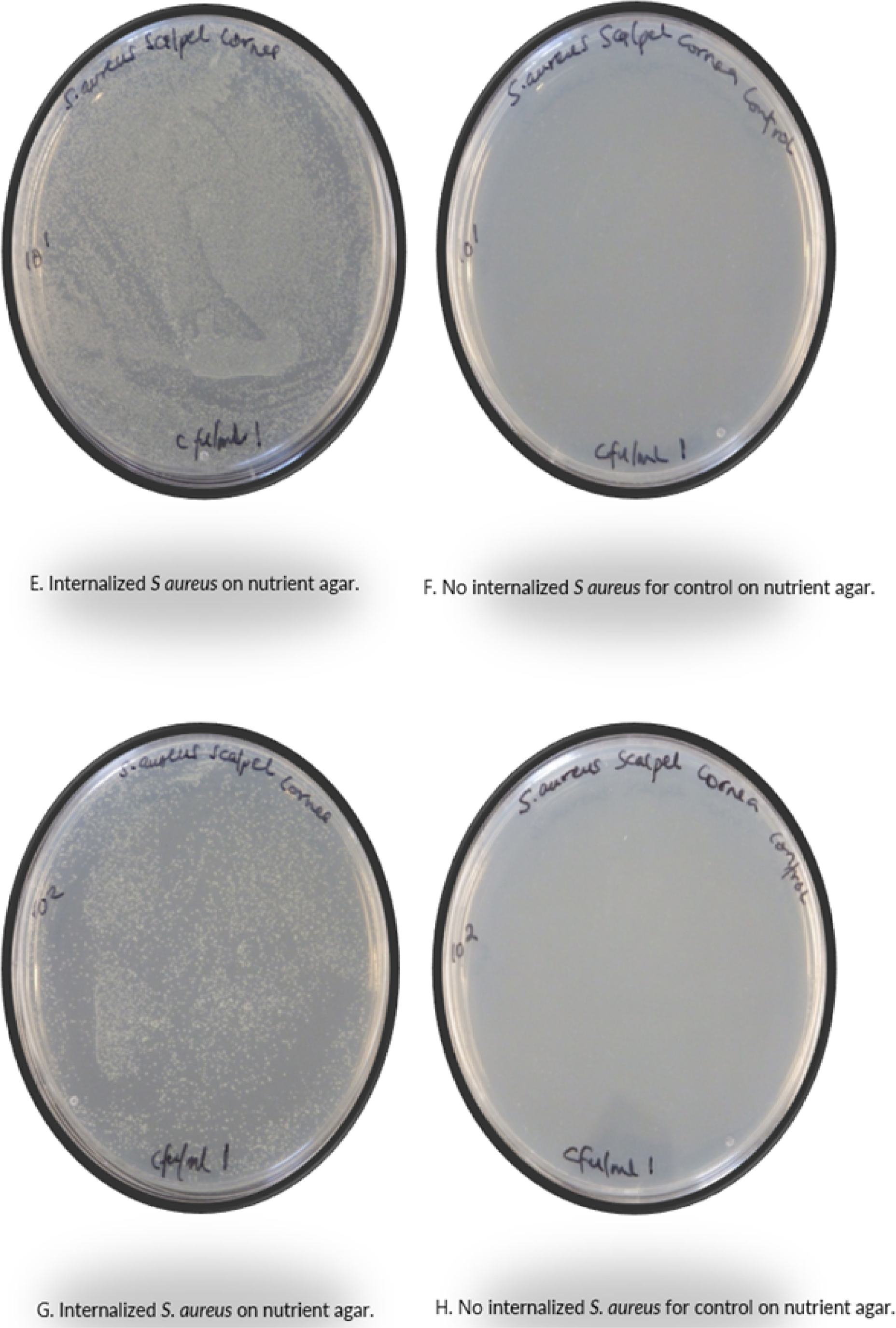
Internalized *S. aureus* from compromised corneas. Excised bovine corneas were compromised by either incision with needles (A, B, C and D) or scarification with scalpels (E, F, G and H). These were inoculated with *S. aureus* and sampled following 1%(w/v) saponin treatment on nutrient agar plates incubated at 37°C for 24 hours. Bacteria were recovered from infected corneas (A, C, E, and G), but not uninfected controls (B, D, F, and H). Quantitative findings obtained from presence or absence of internalized bacteria for serial dilutions of up to 10^2^.

**Figure 7:**
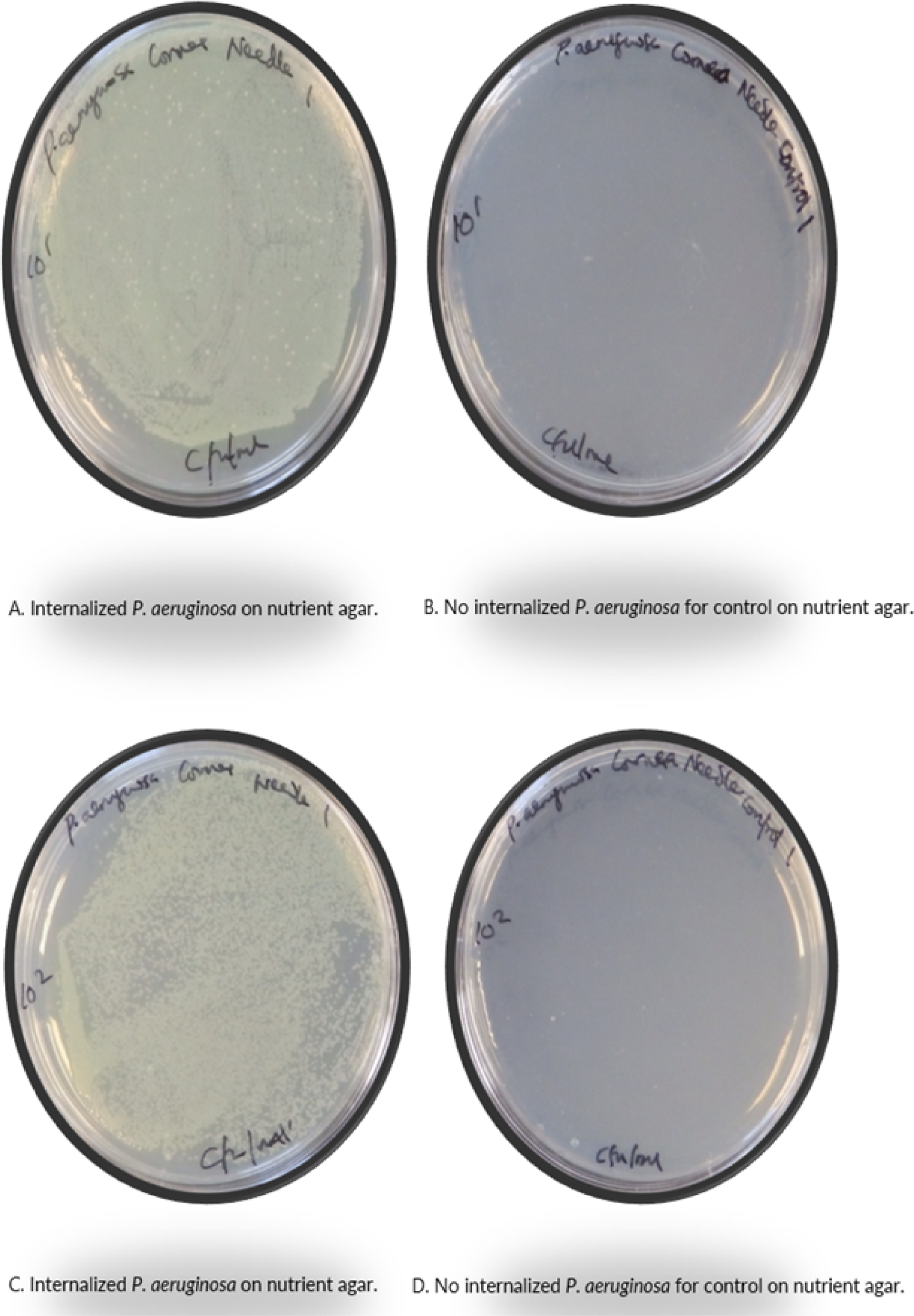
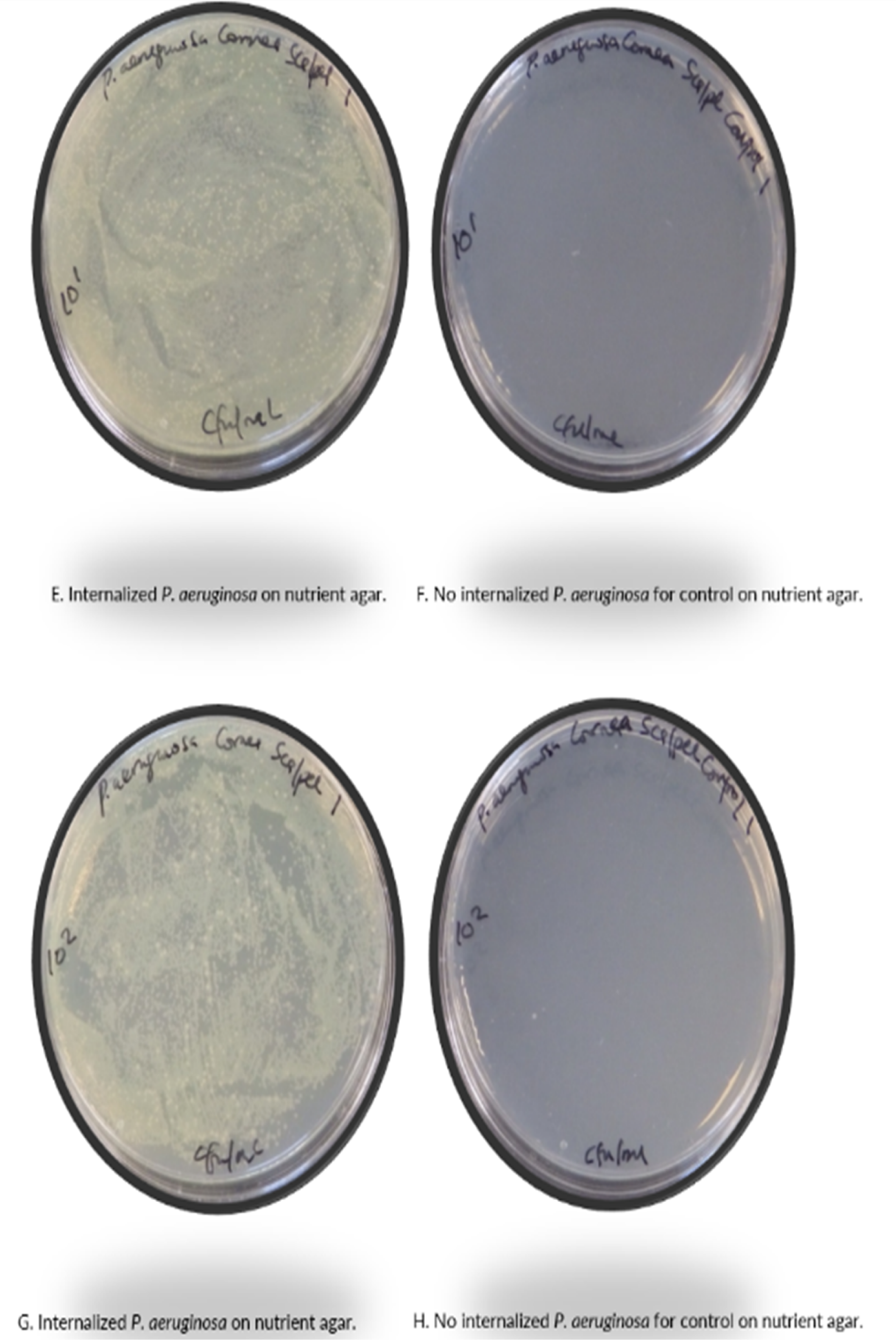
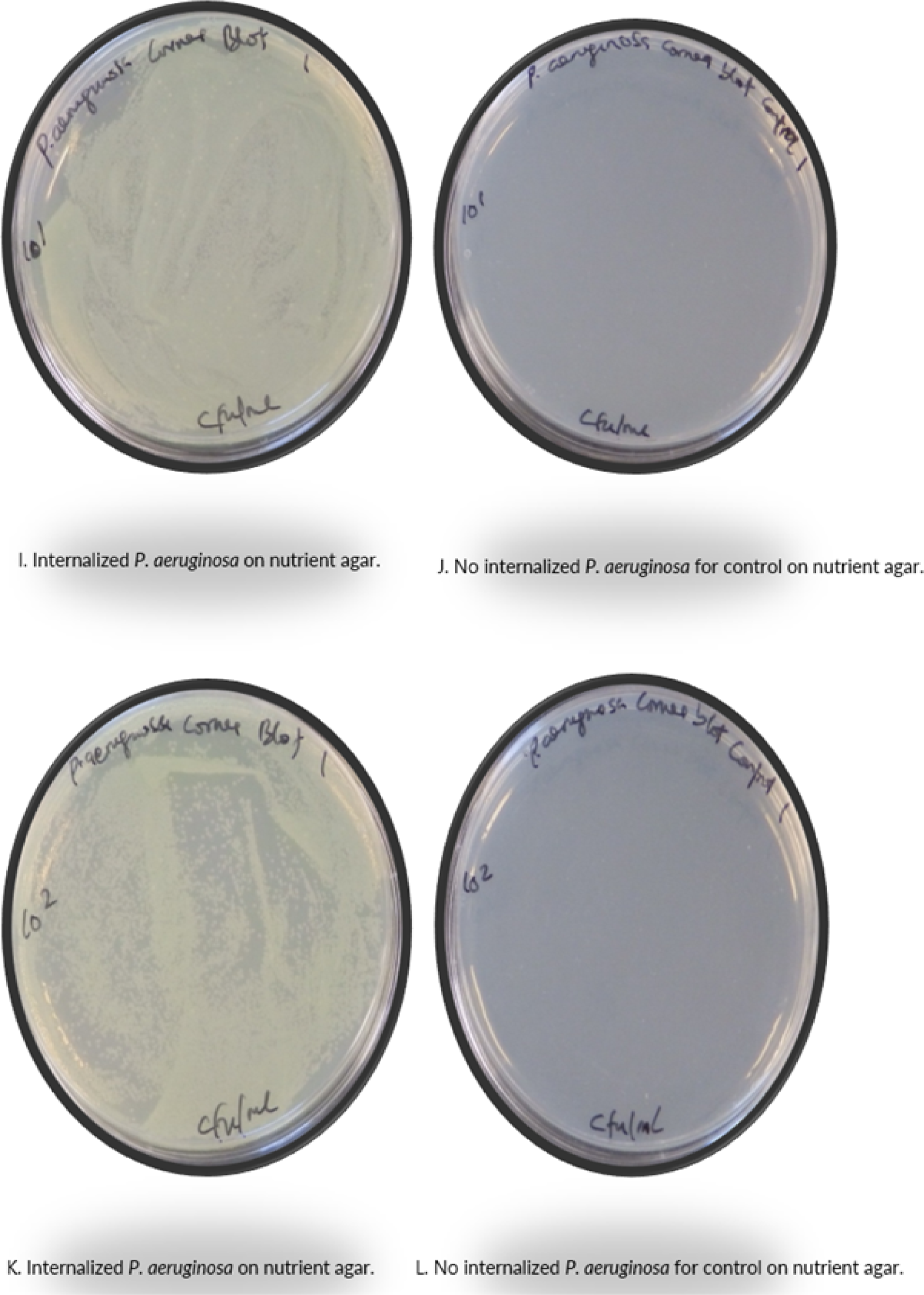
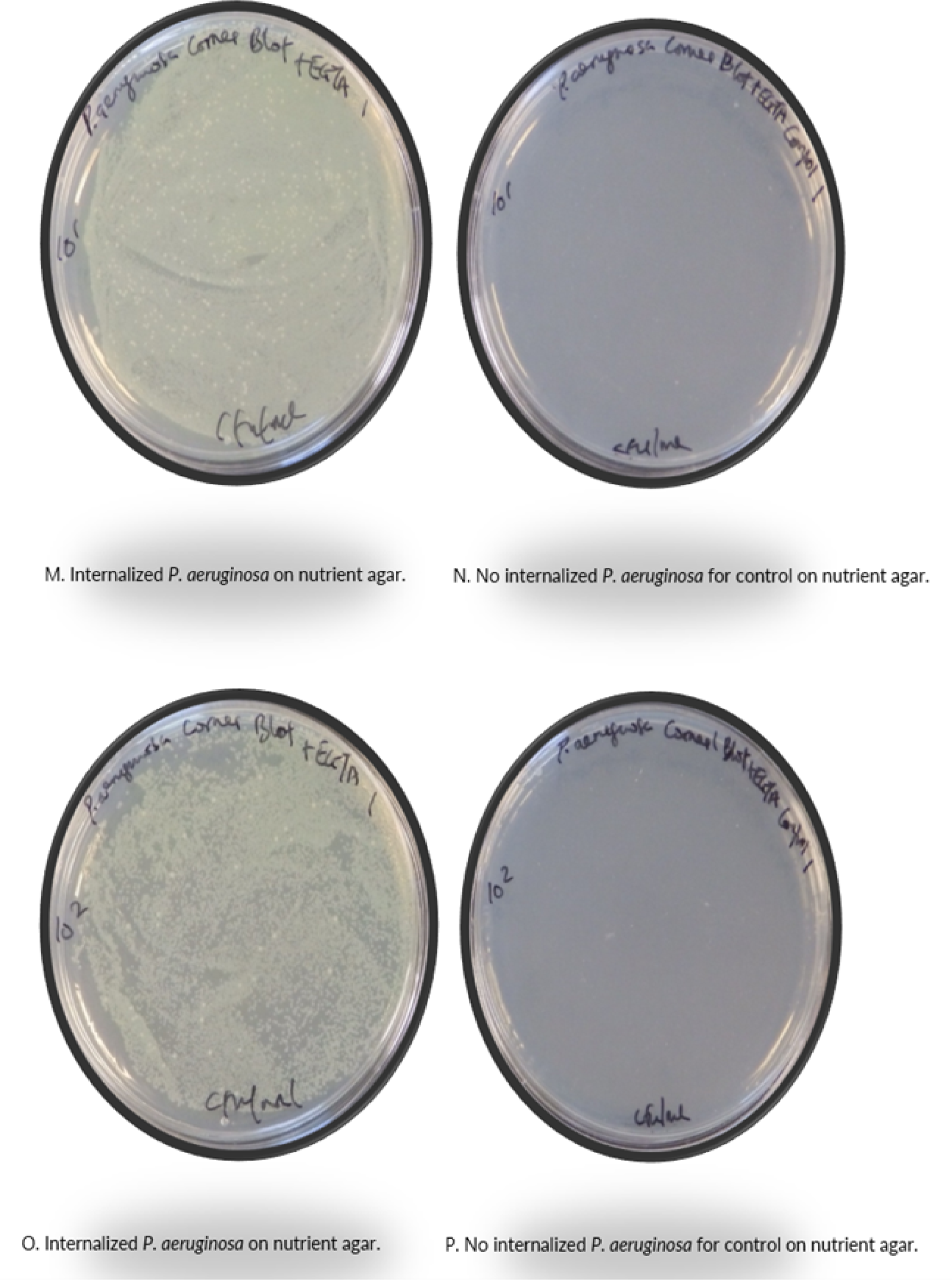
Internalized *P. aeruginosa* recovered from corneas with compromised epithelium. Excised bovine corneas were compromised with needles (A, B, C, and D), use of scalpels (E, F, G, and H), blotting with paper towel (I, J, K, and L), or blotting + treatment with EGTA (M, N, O, and P). Infection with *P. aeruginosa* and internalization was assessed using 1% (w/v) saponin treatment, sampled on nutrient agar plates incubated for 24 hours at 37°C. There was recovery of *P. Aeruginosa* from compromised, infected corneas (A, C, E, G, I, K, M, and O), not from the controls (B, D, F, H, J, L, N, and P). Quantitative data is reported as presence or absence of internalized bacteria for serial dilutions of up to 10^2^.

### 3.2 Antimicrobial susceptibility of bacteria to antibiotics

The MIC and MBC of ceftriaxone against *N. gonorrhoeae* was 0.061035 ± 0.030518 µg/ml and 0.12207 ± 0.066706 µg/ml, respectively. The MIC of ciprofloxacin was lower for *S. aureus* (0.015259 ± 0.008624 µg/ml) than *P. aeruginosa* (0.061035 ± 0.040371 µg/ml), while the MBC was higher for *S. aureus* (0.24414 ± 0 µg/ml) than *P. aeruginosa* (0.12207 ± 0.061035 µg/ml) (Fig 8 and Table 1).

**Fig 8:**
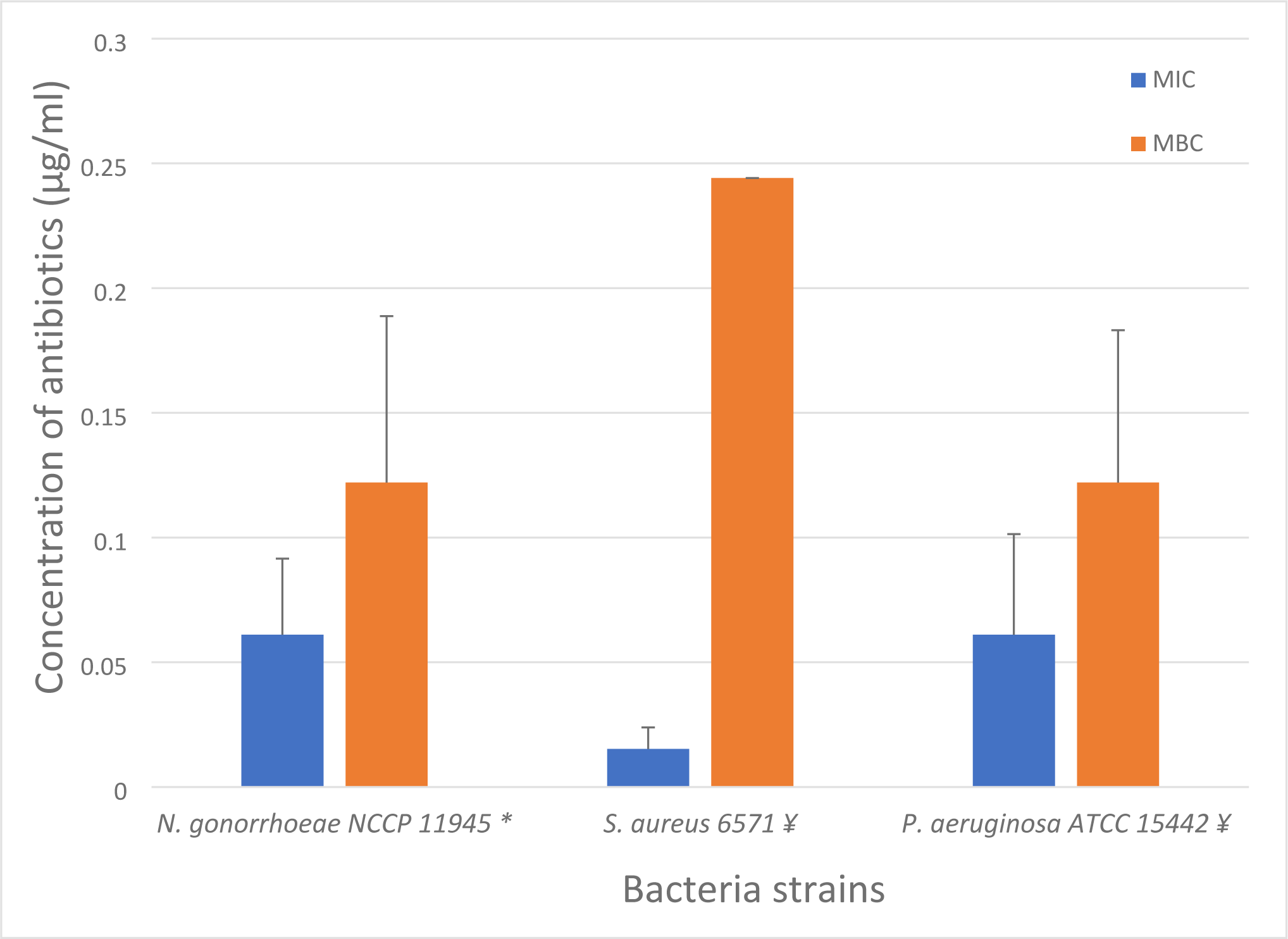
Antibacterial activity of ceftriaxone against *N. gonorrhoeae*, and ciprofloxacin against *S. aureus and P. aeruginosa.* Ceftriaxone challenged *N. gonorrhoeae* (∼ 10^6^ cfu/ml), and ciprofloxacin challenged *S. aureus* and *P. aeruginosa* (∼ 10^5^ cfu/ml) to determine the minimum inhibitory concentrations (MICs) denoted as the least concentrations of the antibiotics that inhibited growth of bacteria after 24 hours. Concentrations ranged from 500 µg/ml to 5.96×10^-5^ µg/ml). The minimum bactericidal concentrations (MBCs) of the antibiotics at which no bacteria growth occurred when spotted from 96 well plates on the respective agar plates (GC agar for *N. gonorrhoeae* and nutrient agar for *S. aureus* and *P. aeruginosa*) following 24 hours of incubation. The data denotes mean result of triplicates for n=3 independent experiments. ᵡ Ceftriaxone was tested against *N. gonorrhoeae* NCCP 11945 ¥ Ciprofloxacin was tested against *P. aeruginosa* ATCC 15442 and *S. aureus* 6571.

**Table 1:** Antimicrobial activity of ceftriaxone against *N. gonorrhoeae* and ciprofloxacin against S. *aureus* and *P. aeruginosa*.

| **Bacteria strains** | **Mean**<br>**MICs\***<br>**(µg/ml)** | **Standard**<br>**deviations**<br>**(STDev)** | **Mean MBCs^**<br>**(µg/ml)** | **Standard**<br>**deviations**<br>**(STDev)** |
| --- | --- | --- | --- | --- |
| ***N. gonorrhoeae***<br>**NCCP 11945** | 0.061035 | 0.030518 | 0.12207 | 0.066706 |
| ***S. aureus* 6571** | 0.015259 | 0.008624 | 0.244141 | 0 |
| ***P. aeruginosa***<br>**ATCC 15442** | 0.061035 | 0.040371 | 0.12207 | 0.061035 |

\* MICs: Minimum inhibitory concentrations.
^ MBCs: Minimum bactericidal concentrations.

### 3.3 Clearance of corneal infection with antibiotics

Corneas infected with *N. gonorrhoeae* were treated afterwards with ceftriaxone and those infected with *S. aureus* or *P. aeruginosa* were treated thereafter with ciprofloxacin for 1 hour. It was observed that no bacterial colonies were recovered from corneas treated with antibiotics in comparison to the PBS controls (Figs 9-11).

**Fig 9:**
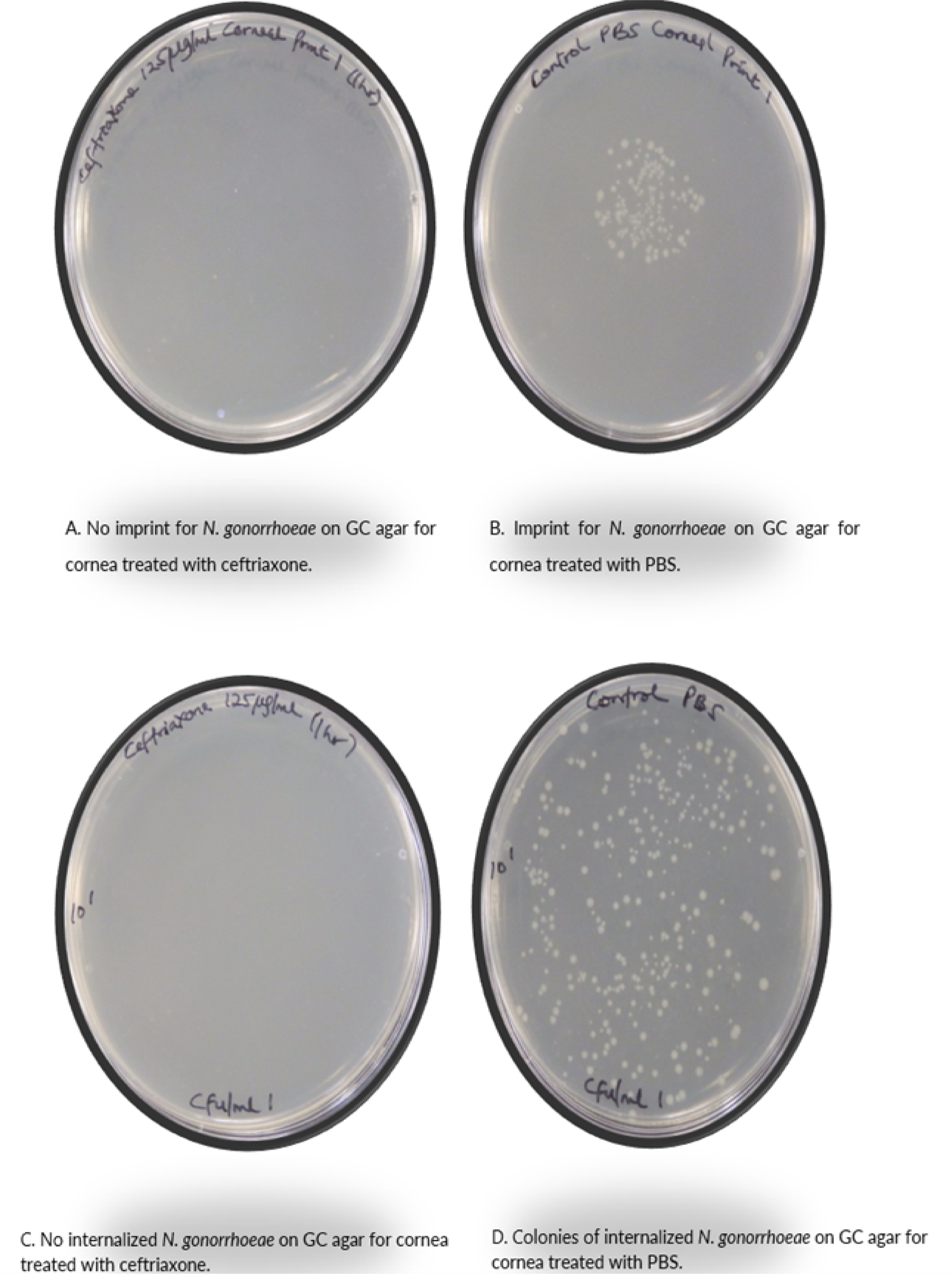
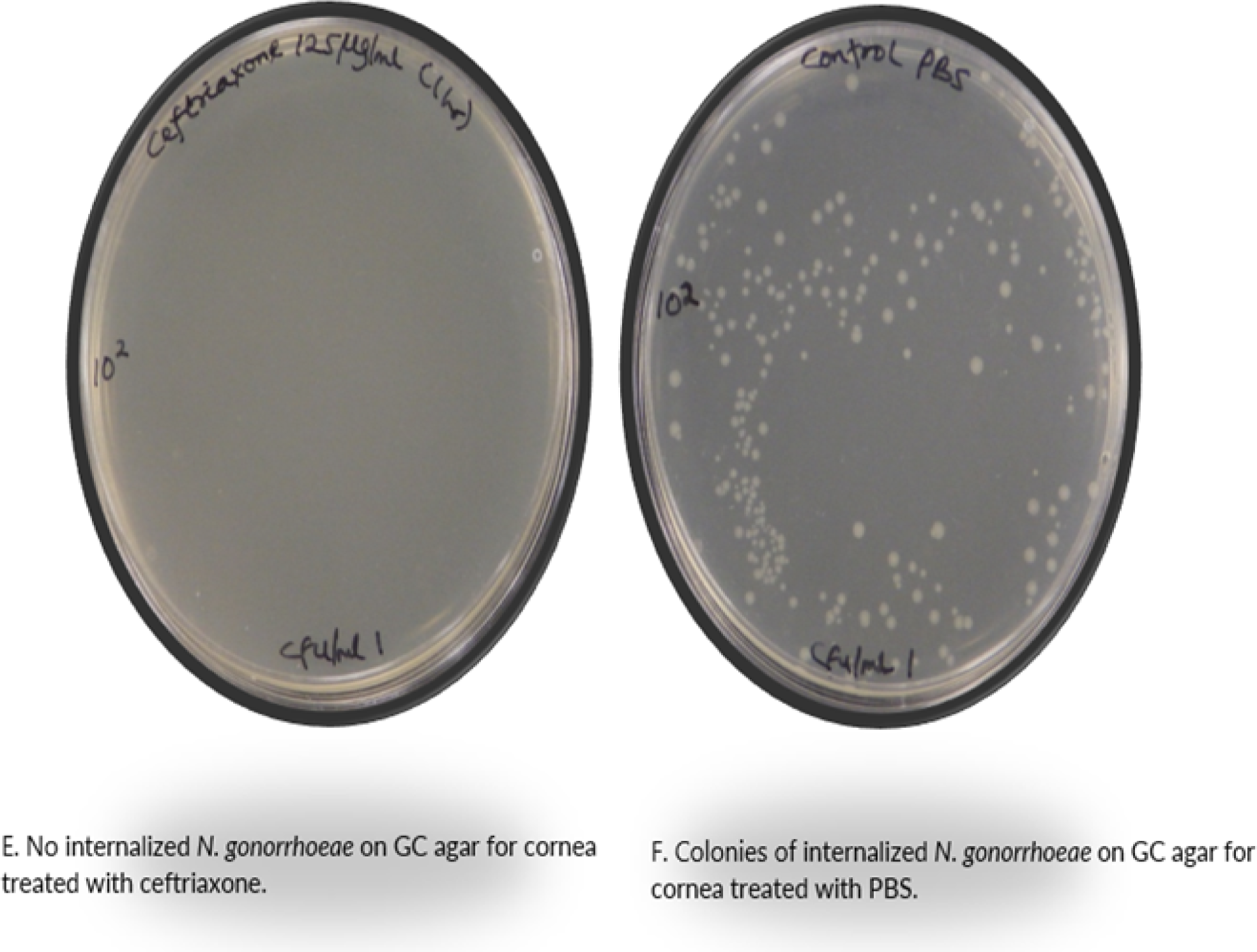
Treatment of gonococcal infected bovine corneas with ceftriaxone. Excised bovine corneas infected with *N. gonorrhoeae* were treated with 125 µg/ml ceftriaxone or PBS as control and sampled on GC agar plates incubated in 5% CO_2_ for 24 hours at 37°C. No bacterial colony was recovered from corneas treated for a duration of 1 hour with ceftriaxone (A, C, and E) in comparison to controls treated with PBS (B, D, F). Results for bacteria adherence are based on corneal imprints (A, B) and for internalized bacteria based on saponin treated serial dilutions (C, D, E, F).

**Fig 10:**
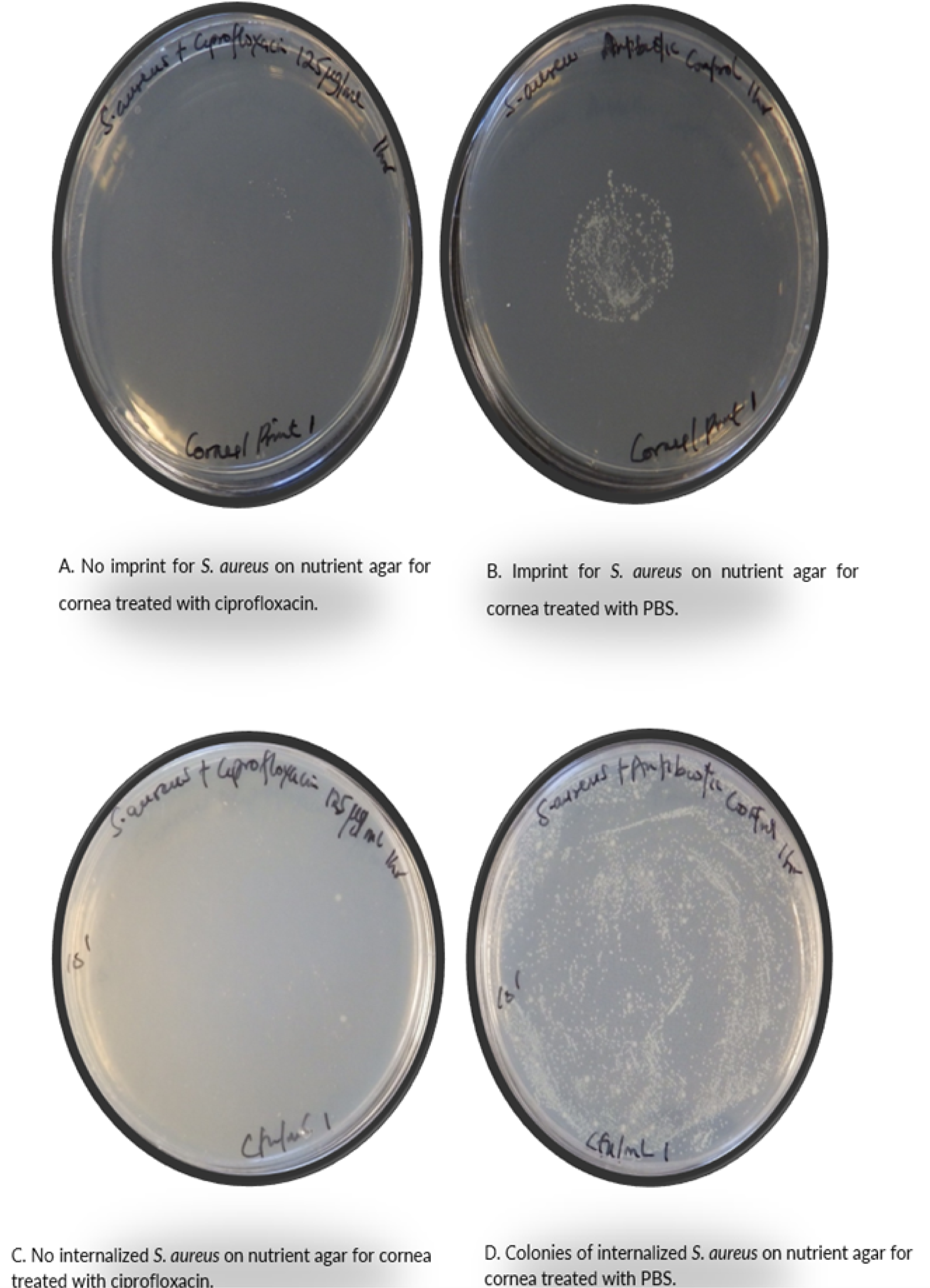
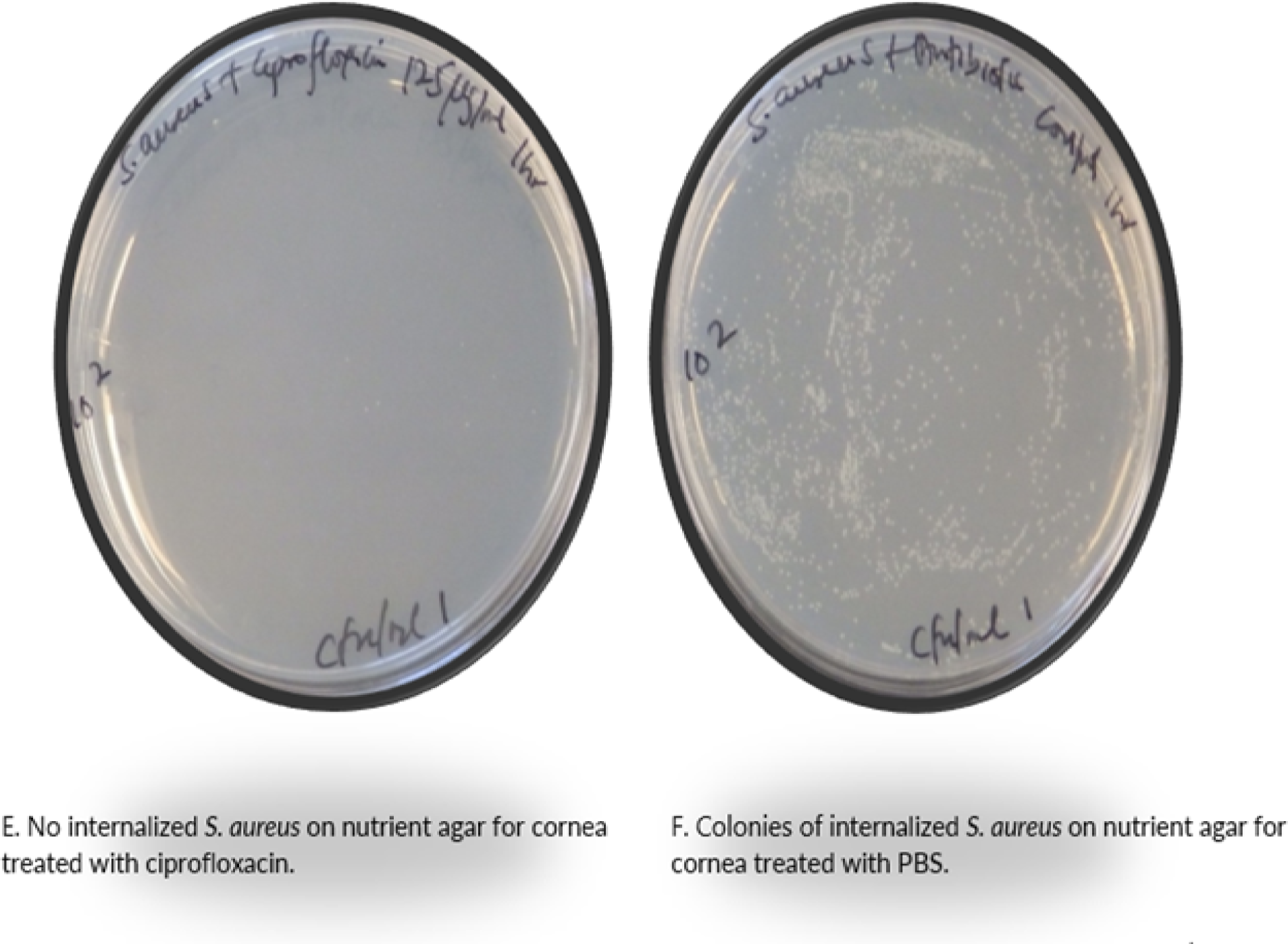
Treatment of staphylococcal infected bovine corneas with ciprofloxacin. Excised corneas of bovine eyes infected with *S. aureus* were treated with 125 µg/ml ciprofloxacin, with PBS as control, and enumerated on nutrient agar plates incubated at 37°C for 24 hours. There was no bacterial growth observed on the nutrient agar plates following ciprofloxacin treatment of corneas for 1 hour (A, C, and E) in contrast to normal growth on controls treated with PBS (B, D, F). Findings for bacteria adherence is based on corneal imprints for the infection experiment (A, B) and serial dilutions of released internalized bacteria (C, D, E, F).

**Fig 11:**
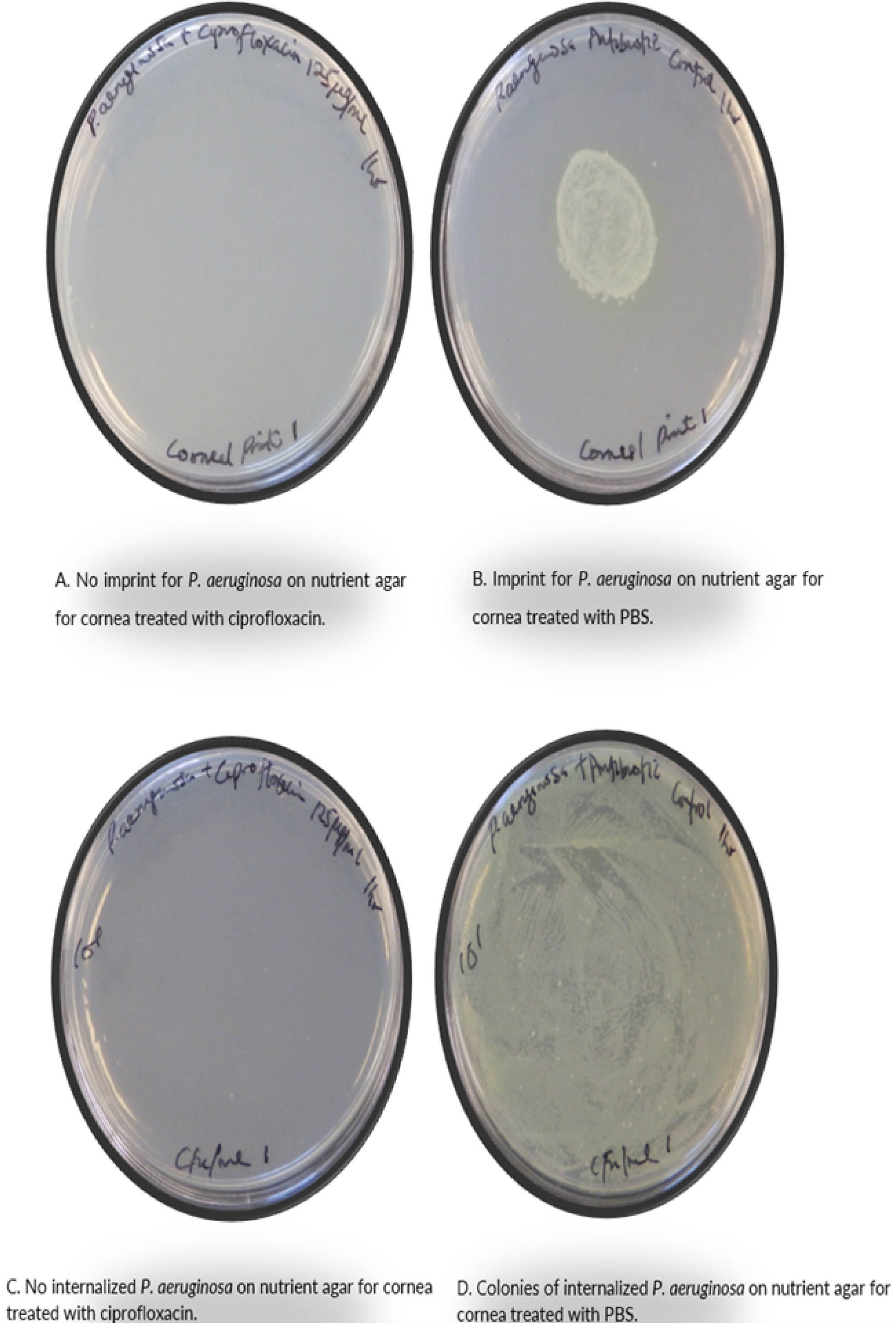
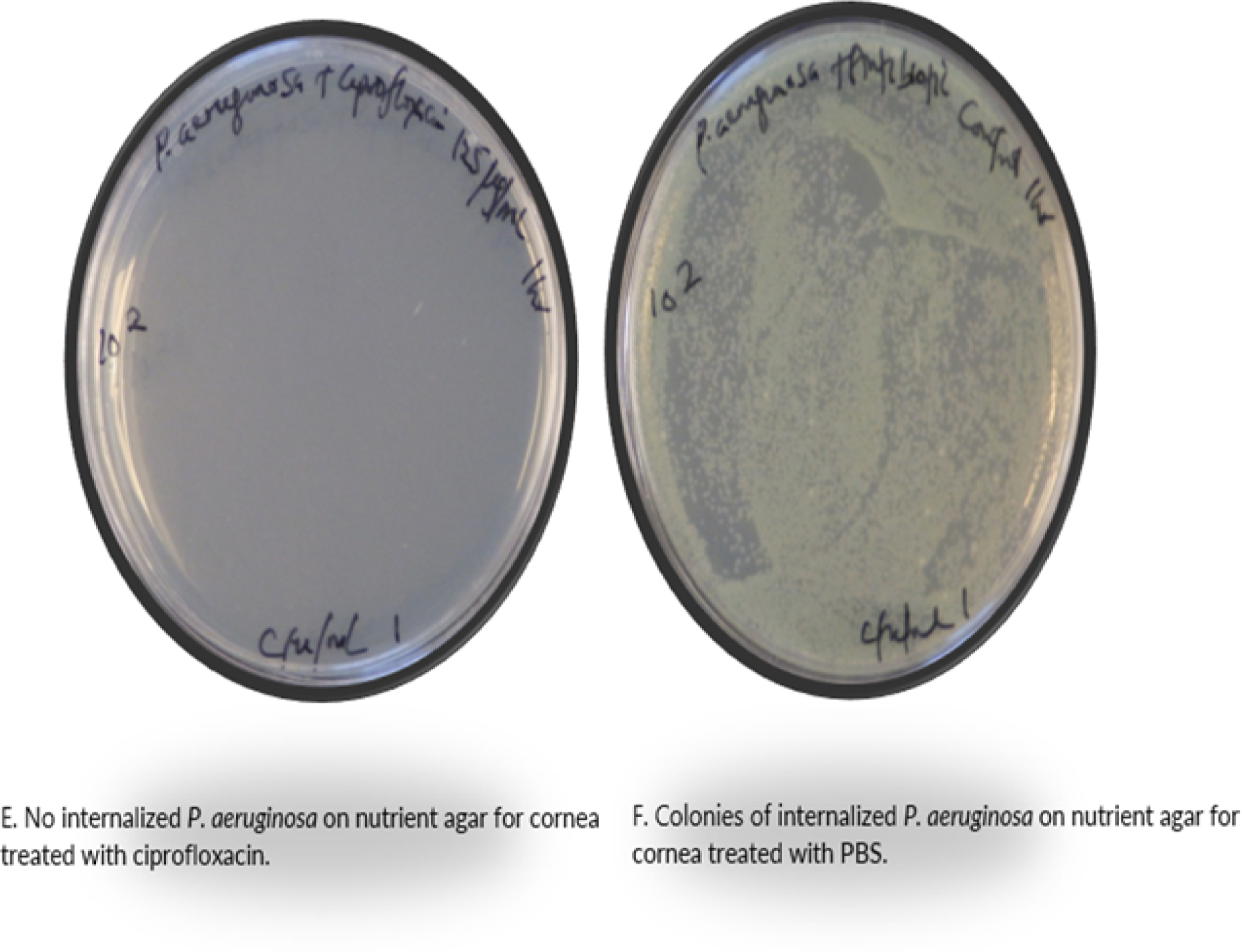
Treatment of pseudomonad infected bovine corneas with ciprofloxacin. Pseudomonad infected excised bovine corneas were challenged with 125 µg/ml ciprofloxacin (PBS control) and sampled on nutrient agar plates incubated for 24 hours at 37°C. No bacterial colonies were found for corneas treated for a duration of 1 hour with ciprofloxacin (A, C and E), unlike controls treated with PBS (B, D, F). Data for bacteria adherence is based on corneal imprints for the infection experiment (A, B) and internalized bacteria serial dilutions (C, D, E, F).

## 4.0 Discussion

An *ex vivo* bovine eye model for investigating corneal bacterial infection and clearance of infection, offers a relatively quick, easy, and cost-effective option. The bovine eyes used in this study were obtained as by-products from an abattoir. This eliminates the time to feed and prepare live animals for experimentation, as well as unnecessary suffering of animals that may occur during experiments in spite of ethical principles that guide the use of animal experiments [51–53,55,86,87, 97–100]. In this *ex vivo* bovine eye model, corneal infection with *N. gonorrhoeae* was established at 1 hour of infection (Figs 3A, 5A, and 5C), *S. aureus* at 4 hours (Figs 3C, 4A, 4C, 5E, 5G, 6A, 6C, 6E, 6G) and by 6 hours *P. aeruginosa* infection was demonstrated (Figs 3E, 4E, 4G, 4I, 4K, 5I, 5K, 7A, 7C, 7E, 7G, 7I, 7K, 7M, and 7O). These results correlate well with clinical manifestations of these bacterial infections, equating to the time necessary *in vivo* to establish infections. This model will provide valuable insight when assessing antimicrobial agents to be used for prophylaxis and treatment in bacterial ocular infections [19, 50]. For instance, it has been suggested that the promptness of administering a prophylactic agent in gonococcal ophthalmia neonatorum can determine whether the intervention would succeed [101]. Due to the potential for *N. gonorrhoeae* infection to rapidly cause ocular complications to the cornea and even blindness [9,43–49], recommendations have been made for administration of prophylaxis to the new-born after birth ranging from within 24 hours [102], within four hours [101], within one hour [103], to immediately after birth [104]. This emphasizes the importance of novel antimicrobials such as monocaprin, which clears the bacteria from the cornea within 2 minutes [105].

The dynamics of inter- and intrastrain bacterial variation and other factors occurring at the corneal surface may impact establishment of infection. *P. aeruginosa* strain Pa01 from three different sources showed variable ability to penetrate the corneal epithelium in a murine *ex vivo* infection model [50]. In another study involving a mouse model, *P. aeruginosa* isolated from human eye infections or inflammation showed different corneal pathologies and not all of these isolates were capable of causing corneal infection in the mice [106]. Additionally, corneal epithelial adhesion of *P. aeruginosa* Pa01 occurred after 4.5 hours in murine β-defensin 3 (mBD-3)-deficient mice, compared to after 7.5 hours in wild-type C57BL/6 mice. This suggests that the loss of mBD-3 rendered the murine corneas more susceptible to bacterial colonisation. Despite colonising the cornea in these mouse model investigations, there was no penetration of the bacteria beyond the corneal epithelium [107].

Differences between mammalian species’ ocular tissues may influence bacterial infection and antimicrobial drug permeation, which is vital to reach internalised bacteria. Several differences have been documented to exist between corneas of species such as rabbits, pigs, cows, and humans. While the thickness of porcine and human corneas appears to be similar (0.5 – 0.7mm) [19], corneas of rabbits are thinner (0.3 – 0.4mm) [108–111], with bovine corneas being the thickest (0.9 – 1.1mm) [19, 112]. Similarly, bowman’s layer is thought to be present in both porcine and human corneas but absent in rabbit and bovine corneas [19, 108]. While the epithelial thickness (30 – 40 µm) and cell layers (5 – 7) of rabbits and humans are identical [19,101], porcine corneas are thicker (50 – 70 µm and 6 – 9 layers), and bovine corneas possess the greatest epithelial thickness (70 – 90 µm) and cell layers (10 – 15 layers) [19, 113]. Moreover, corneal length also denoted as corneal horizontal diameter was reported to be lowest for humans (11.7 mm) [19,108,114], followed by pigs (12.1 – 14.2 mm) [19]. Rabbit corneal lengths (15 mm) [19,108] were greater than those of pigs, but less than those of cows (29.3 mm) [19, 112]. Despite similarities in epithelial thickness and cell layers of corneas between rabbits and humans, a study reported recovery of staphylococcal and pseudomonad bacterial colonies from corneas of humans that was tenfold greater than those recovered from rabbits after 24 and 48 hours, post-inoculation [20]. In addition, an inversely proportional relationship is suggested to exist between thickness and membrane transfer rate [115]. A study that investigated permeability of ophthalmic drugs like ciprofloxacin hydrochloride, timolol maleate, and lidocaine hydrochloride observed that rabbit corneas had the highest drug permeability, which was followed by porcine corneas, and the least permeable were bovine corneas [103]. Likely contributors to higher drug permeability in rabbits [116] include a relatively large corneal horizontal diameter (15mm) relative to the horizontal diameter for the globe (18 – 20mm), providing a large surface area for drug absorption and the absence of bowman’s membrane [108]. Comparison of permeation of pharmaceutical formulations such as aceclofenac [117] and gatifloxacin antibiotic [118] showed that maximum drug release occurred in corneas of goats, followed by sheep, and least of those investigated was buffalo. Since a longer path of diffusion may characterise thicker corneas, differences in cornea thickness for these species appeared to account for these observed findings. Corneas of goats are thinner (0.68±0.0003 mm) in comparison to sheep (0.86±0.0003 mm), with those of buffalo being the thickest (1.1±0.0006 mm). This agrees with the highest permeation being seen with caprine corneas [117]. Factors such as these and the influence of transport and binding proteins in the corneal layers should be carefully considered when designing animal experiments including *ex vivo* animal models [116].

Bacterial inoculation of the bovine corneas investigated here involved intact corneal epithelium and epithelium compromised with needle, scalpel, blotting, and blotting with EGTA. Bacteria adhered to the corneal surface and were internalised by compromised as well as uncompromised epithelium. Recovery of *N. gonorrhoeae, S. aureus*, and *P. aeruginosa* from imprints on agar plates showed bacteria adhered to the surface of the infected corneas (Figs 3 and 4), providing qualitative data.

Internalised bacteria were also recovered from the infected corneas (Figs 5-7). Calculation of colony forming units per ml (cfu/ml) provides quantitative data (Fig 12 and Table 2), which shows the activity of the antimicrobial agent in clearing infection from the *ex vivo* bovine corneal model, compared to controls (Figs 9-11).

**Fig 12:**
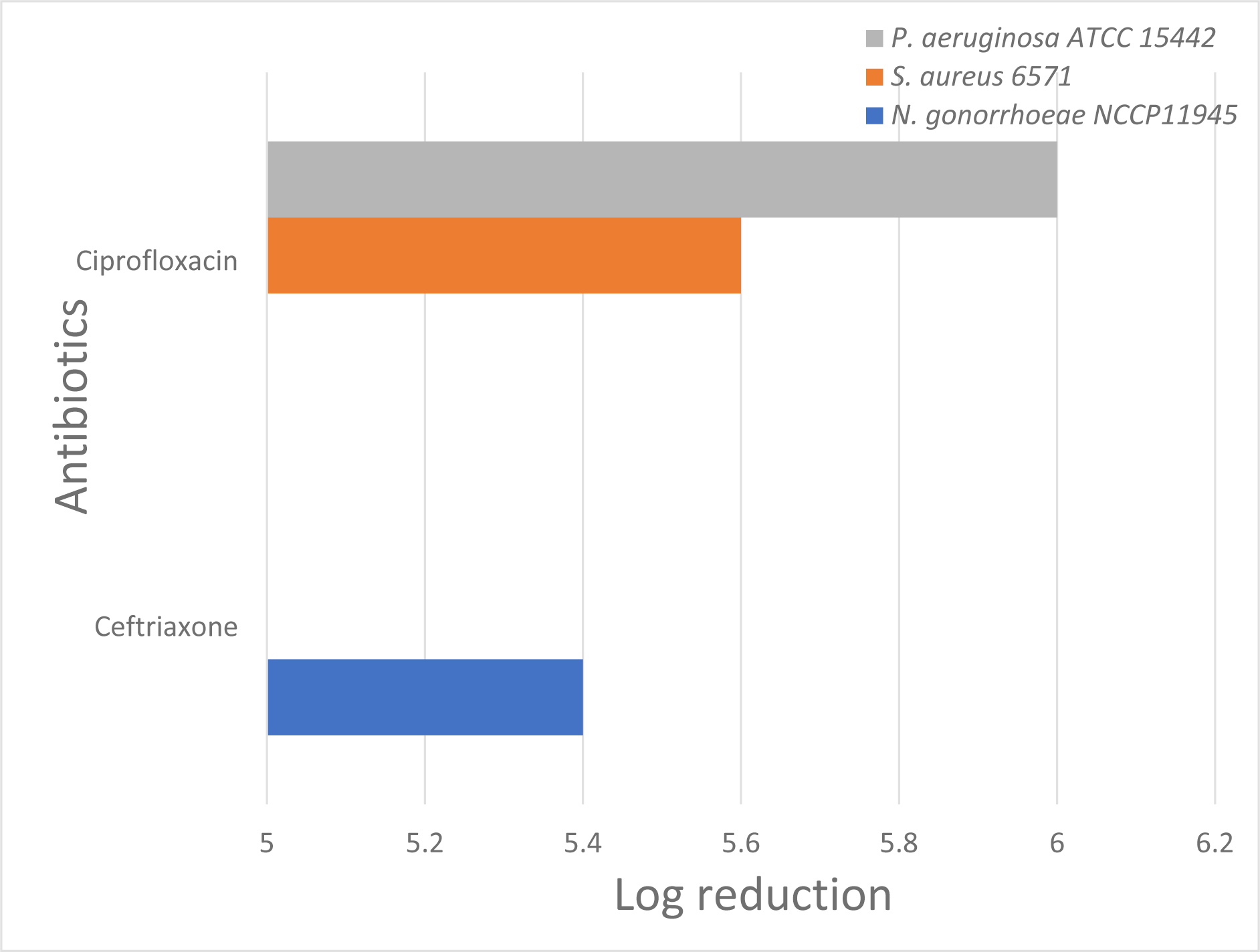
Log reduction of gonococcal, staphylococcal, and pseudomonad clearance from infected bovine corneas. Ceftriaxone (for *N. gonorrhoeae*) and ciprofloxacin (for *S. aureus* and *P. aeruginosa*) at 125 µg/ml were effective at clearing bacteria from infected corneas. A 6 log_10_ reduction was seen for ciprofloxacin on corneas infected with *P. aeruginosa*, 5.6 log_10_ for ciprofloxacin on *S. aureus*, and 5.4 log_10_ for ceftriaxone on *N. gonorrhoeae*. These numbers reflect no growth of bacteria versus growth on the PBS control and are limited by the experimental parameters.

**Table 2:** Bactericidal activity of ceftriaxone and ciprofloxacin in clearance of gonococcal, staphylococcal and pseudomonad infections from excised bovine corneas.

| Concentration of antibiotics | *N. gonorrhoeae*<br>NCCP 11945 | *S. aureus* 6571 | *P. aeruginosa*<br>ATCC 15442 |
| --- | --- | --- | --- |
| Ceftriaxone<br>125 µg/ml | 5.4 | N.D* | N.D* |
| Ciprofloxacin<br>125 µg/ml | N.D* | 5.6 | 6 |

\* N.D: Not done

Furthermore, the results obtained for the infected corneas with intact epithelium and compromised epithelium were comparable (Figs 5E and 5G vs 6A and 6C, 6E and 6G; and Figs 5I and 5K vs 7A and 7C, 7E and 7G, 7I and 7K, 7M and 7O). This shows that after attachment of the bacteria to the corneal epithelium, the bacteria traversed the layers of the cornea beyond the epithelium. There is therefore both cellular attachment and internalisation of the bacteria in this *ex vivo* bovine corneal model.

Bacterial traversal within the cornea has been demonstrated by histological analysis previously [18,20]. Recent advances that do not require histological processing of corneal tissues have enabled the localisation of internalised bacteria within the cornea using specialized microscopy and imaging techniques. Translocation of pseudomonad bacteria has been shown in *ex vivo* mouse models in which corneas were intact and in those compromised (with tissue paper + EGTA treatment). This work demonstrated the roles of factors such as ExSA regulated type III secretion (T3SS), MyD88-mediated signalling, and dendritic cells in regulating this traversal [50,121,122]. Future use of these microscopy techniques might therefore be useful in combination with the *ex vivo* bovine model to provide additional insights into mechanisms associated with corneal infection.

Traversal of *P. aeruginosa* has been reported to occur in immortalised rabbit corneal epithelial cells [22,119] and in multi-layered human corneal epithelial cells cultured *in vitro* [120, 107]. *P. aeruginosa* infected the excised bovine corneas by 6 hours (Figure 7) and was able to internalise into the tissues. Previously, *in vivo* and *ex vivo* mouse models of *P. aeruginosa* infection found the bacteria adhered to the blotted corneas by 5 hours but did not penetrate the corneal epithelium until after 3 hours later [26].

Despite reports in the literature involving other strains [123–128], ceftriaxone was found to be effective against the *N. gonorrhoeae* strain NCCP11945 used here (Fig 8 and Table 1). Based on this data, 125 µg/ml of ceftriaxone was used here. Although resistance has been reported for some strains of *P. aeruginosa* [129–134] and *S. aureus* [135–139] ciprofloxacin was an effective antimicrobial against the strains investigated here based on MICs (Fig 8). Based on these susceptibilities, 125 µg/ml of ciprofloxacin was used to clear these induced infections from the *ex vivo* bovine corneal model. After 1 hour, the corneas were completely clear of bacteria, compared to the negative control (Figs 9-11). The log reduction can therefore be calculated to quantitatively show the antibiotic killing of these species of bacteria infecting the corneas and hence the bactericidal activity in the infected *ex vivo* bovine corneal model just like in *in vivo* models [27,33,140, 141]. The bactericidal activity of the antibiotics at 125 µg/ml that cleared the bacterial infections from the corneas (Figs 9-11) was at least 5 log_10_ reduction (Fig 12 and Table 2). This value is restricted due to the design of the experiment and is calculated based on the absence of bacterial growth after treatment, observed on the respective agar plates, in comparison to the negative control (PBS).

Conventionally, erythromycin ointment has been utilised for prophylaxis / treatment of neonatal gonococcal conjunctivitis [104,142,143]. Ceftriaxone on the other hand is given intramuscularly for treatment of *N. gonorrhoeae* conjunctivitis in new-borns [144,145] and adults [146,147]. However, it is imperative that more topical options for prophylaxis and treatment of neonatal gonococcal conjunctivitis are investigated.

Ceftriaxone cleared *N. gonorrhoeae* from infected bovine corneas within 1 hour. An initial assessment would suggest that there is potential for consideration of this antibiotic for neonatal gonococcal conjunctivitis and perhaps other microbial conjunctivitis including those seen in adults by way of re-purposing this antibiotic for topical application. However, ceftriaxone precipitates in solutions containing calcium, making it unsuitable for use on the eye due to the presence of calcium in human tear fluid. Whilst there could be pharmaceutical formulations able to chelate out calcium, this may be physiologically harmful to the eye. In the case of new-borns up to 28 days old, ceftriaxone is contraindicated due to the potential need for calcium treatment, which can be required in premature infants and those with impaired bilirubin binding [148–151]. Thus, although further investigations may be able to circumvent these issues with ceftriaxone, alternatives are needed.

*P. aeruginosa* and *S. aureus* are also implicated in the aetiology of ophthalmia neonatorum [152–157], as well as microbial keratitis in adults including in cases of contact lens wear [7,158–166]. Fluoroquinolones have also been used for treatment of ophthalmia neonatorum [143] and microbial keratitis [166–169]. According to The American Academy of Paediatrics, 0.5% erythromycin should be given as ocular prophylaxis to neonates. If this is not available, 1% azithromycin is suggested with 0.3% ciprofloxacin ophthalmic ointment being suggested as a potential in the absence of both erythromycin and azithromycin topicals [170]. Ciprofloxacin was shown here to be effective in clearing staphylococcal and pseudomonad infections at 125 µg/ml, which is less than the 3,000 µg/ml in the 0.3% ciprofloxacin ophthalmic formulation [171].

Ocular defences enable the cornea to resist invasion by microorganisms. For example, *in vivo* infection of healthy animal corneas with *P. aeruginosa* did not result in inflammation and infection due to bacterial clearance by the ocular defences [172]. These include the epithelial barrier of the cornea, tear exchange and eye blinks, and upregulation or expression of biochemical components such as defensin, secretory immunoglobulin A, mucin glycoproteins, antimicrobial proteins, and peptides [173–176]. Innate defences of surfactant proteins A and D (SP-A and SP-D) present on the ocular surface [177], for instance, have been reported to prevent invasion of corneal epithelial cells by bacteria pathogens such as *P. aeruginosa* and *S. aureus* [172,178,179]. Although several immune responses of the ocular system are thought to exist in *in vivo* models, in *ex vivo* models the cellular and stromal constituents of the corneal tissue remain, as well as certain inherent intracellular immune elements [20]. Activity from *ex vivo* tissue was demonstrated from scrapings of human cadaver corneal epithelium. When introduced into organ culture, the scrapings induced epithelial regeneration, demonstrated by an upregulation of the antimicrobial defensin Human ß-defensins 1 and 2 (hBD-1, -2) that promoted re-epithelialization [174], indicating that some of the innate defences may still be present in dead corneal tissues. Further investigation of the *ex vivo* bovine eye model may show it to be comparable to *in vivo* models, despite the potential scarcity or lack of physiological mechanisms of defence.

The defence mechanism provided by the presence of tear fluid and its antimicrobial constituents has also been argued in favour of *in vivo* models [180,181]. However, alternative study designs that may add artificial tears or other immune factors to the *ex vivo* bovine eye model may prove useful [182]. In one study, antimicrobial fatty acids and monoglycerides such as myristoleic acid and monocaprin were found to be effective against *N. gonorrhoeae in vitro* and in artificial tears while other anti-gonococcal fatty acids were less potent against *N. gonorrhoeae* in artificial tears [8]. This suggests that study designs other than *in vivo* models can be useful to understand the role of tear fluid as a physiologic mechanism of defence in ocular eye infection models. Furthermore, the biocompatibility of the antibiotics or any selected antimicrobial agent used to clear infection from the cornea in the *ex vivo* bovine eye model can be investigated by histologic assessment of the layers of the corneal tissues and this result may also be comparable with *in vivo* models.

This *ex vivo* bovine eye model provides flexibility of achieving corneal infection in a variety of conditions, thus serving as a good model for simulating conditions of infections that occur with intact corneal epithelium such as in ophthalmia neonatorum and with compromised epithelium seen during contact lens wear. The *ex vivo* approach is also useful in investigations where the experimental design may require exclusion of the impact of tear fluid and certain antimicrobial peptides which inhibit bacteria adhesion, traversal, and viability [95]. The demonstration of corneal infection using this *ex vivo* bovine model with *N. gonorrhoeae, S. aureus,* and *P. aeruginosa* infections suggest that there is the potential for this model to work for other species of microorganisms. Similar *ex vivo* models have been used previously to investigate protozoa [183], viruses [184], and fungi [18,20,183,185]. The usefulness of the bovine *ex vivo* model is demonstrated here and based on this proof of principle, it may be possible to also use porcine [182,185], ovine, or even caprine eyes [18,117]. Future investigations may be needed to compare corneal infection and clearance of infection with the *ex vivo* bovine eye model against *in vivo* animal eye models. Similarly, histological assessment of the infected corneal tissues of the bovine eye model may be investigated to determine the extent that bacteria traverse the different corneal layers and the biocompatibility of an antimicrobial ophthalmic formulation on the cornea in comparison to current *in vivo* eye models. Deciding the appropriate animal model that suits an experiment and harmonising the various available *ex vivo* methods in terms of bacteria strains, inoculation size, infection methods, and incubation times [186] with a view to achieving standardisation would help to enable interstudy comparison and advance the 3R doctrine.

## 5.0 Conclusion

This *ex vivo* bovine eye model is a viable means to study the fundamental processes that characterise bacterial pathogenicity in corneal tissues. In addition, it provides a model system to investigate the effectiveness of conventional and novel ophthalmic formulations against bacterial ocular infections. This model satisfies the need for novel eye models that can investigate a variety of bacterial eye infections caused by strains of *N. gonorrhoeae, S. aureus*, and *P. aeruginosa*, and perhaps other species. As demonstrated here, the model supports experiments that include infection of intact and compromised corneal epithelium. The bovine eyes, and perhaps porcine, ovine, and caprine, can be obtained from the existing human food supply chain, limiting the need for further exploitation of animals in laboratory experiments.

This *ex vivo* corneal model also offers the advantage of being cost effective and time saving. There is no need to further culture corneas in any growth media before they are infected with bacteria. The *ex vivo* bovine corneal infection model shows promise as a viable link between *in vitro* and *in vivo* models and, if optimised, may serve as an alternative to currently existing *in vivo* animal models of corneal infection.

## Declarations

### Data Availability

All relevant data are within the manuscript and supporting files.

